# A trio-binning approach for Cannabis genome *de novo* assembly reveals extensive structural variation, and defines paralog cohorts with very good resolution

**DOI:** 10.1101/2025.05.04.652121

**Authors:** Brett Pike, Alex Kozik, Wilson Terán

## Abstract

With the advent of long read DNA sequencing technologies, assembling eukaryotic genomes has become routine; however, properly phasing the maternal and paternal contributions remains technically challenging. Here, we use the trio-binning approach to separate Oxford Nanopore reads derived from a Cannabis F1 wide cross, made between the Colombian landrace Punto Rojo and the Colorado CBD clone Cherry Pie #16. Reads were obtained from a single PromethION flow cell, generating assemblies with coverage of just 18x per haplotype, but with good contiguity and gene completeness, demonstrating that it is a cost-effective approach for genome-wide and high-quality haplotype phasing, which is of great value for crop breeding programs. Evaluated through the lenses of disease resistance and secondary metabolite synthesis, both being traits of interest for the Cannabis industry, we report copy number and structural variation that, as has recently been shown for other major crops, may contribute to phenotypic variation along several relevant dimensions.

## Introduction

Following 100 years of prohibition, Cannabis is again legal in many countries and jurisdictions. Given its economic and cultural importance, it is notable that genetic resources are scant^1^.

Cannabis is a dioecious annual crop, and its closest relative is *Humulus*, a genera of three species whose most famous member is *H. lupus*, or brewers’ hops. Divergence from their common ancestor is thought to have taken place about 28 MYA in what is today northeast Tibet^2^. Landraces spread to Southeast and Southwest Asia^3^, and later, among other dispersals, to Africa and then South America^4^.

As a natural outcrosser^5^, Cannabis has a low tolerance for inbreeding^6^, and so inbred lines are virtually unknown^7^. The quality of available seedlots was and is highly variable^7^, and so the large majority of production continues to be based on asexual propagation, because clones flower more readily and offer consistent yields of known chemotypes. However, this labor intensive process accounts for as much as 40% of current production costs, reduces genetic diversity, and encourages the persistence of viruses and fungi. Production of uniform, stable lots of F_1_ seed could create economic efficiency and interrupt the disease cycle, with benefits for producers and consumers alike.

In many crops, marker-assisted selection has accelerated the development of new improved lines^8^. Finding markers that flank QTL, and subsequently identifying candidate genes, is facilitated by mapping to a high quality reference genome. The first Cannabis genome to be anointed as the reference by NCBI^9^, a CBD type from Colorado called cs10, offers good contiguity and genic content, and so we have used it as the primary point of comparison in our analyses. However, as a collapsed pseudohaploid, its scaffolds cannot represent the true range of variation found within an individual, and as a modern polyhybrid, it cannot inform as to the ancestral state of the Cannabis population’s founders. In an effort to address this lacuna, we have sequenced both parental haplotypes of an F_1_ derived from two distantly related parents, Punto Rojo (PR) and Cherry Pie (CP), which vary for several important traits of interest.

Punto Rojo is thought to descend from dual-use (drug and fibre) African cannabis introduced to Colombia in the 17th century^4^, and has acclimatized almost entirely in the absence of irrigation, fertilization, and agrochemicals. It has good resistance to fungi and grows well in high heat and low-nutrient soil. The name translates as ‘Red Point’ and refers to the unusual levels of anthocyanin sometimes seen in new shoots and receptive calyces (Fig. 1). In the 60s and 70s, illicit shipments of Type I (THC-dominant) Punto Rojo found favor among American consumers due to its special effects, which were thought to be more psychedelic and less soporific than other imports^10^.

**Figure 1.**
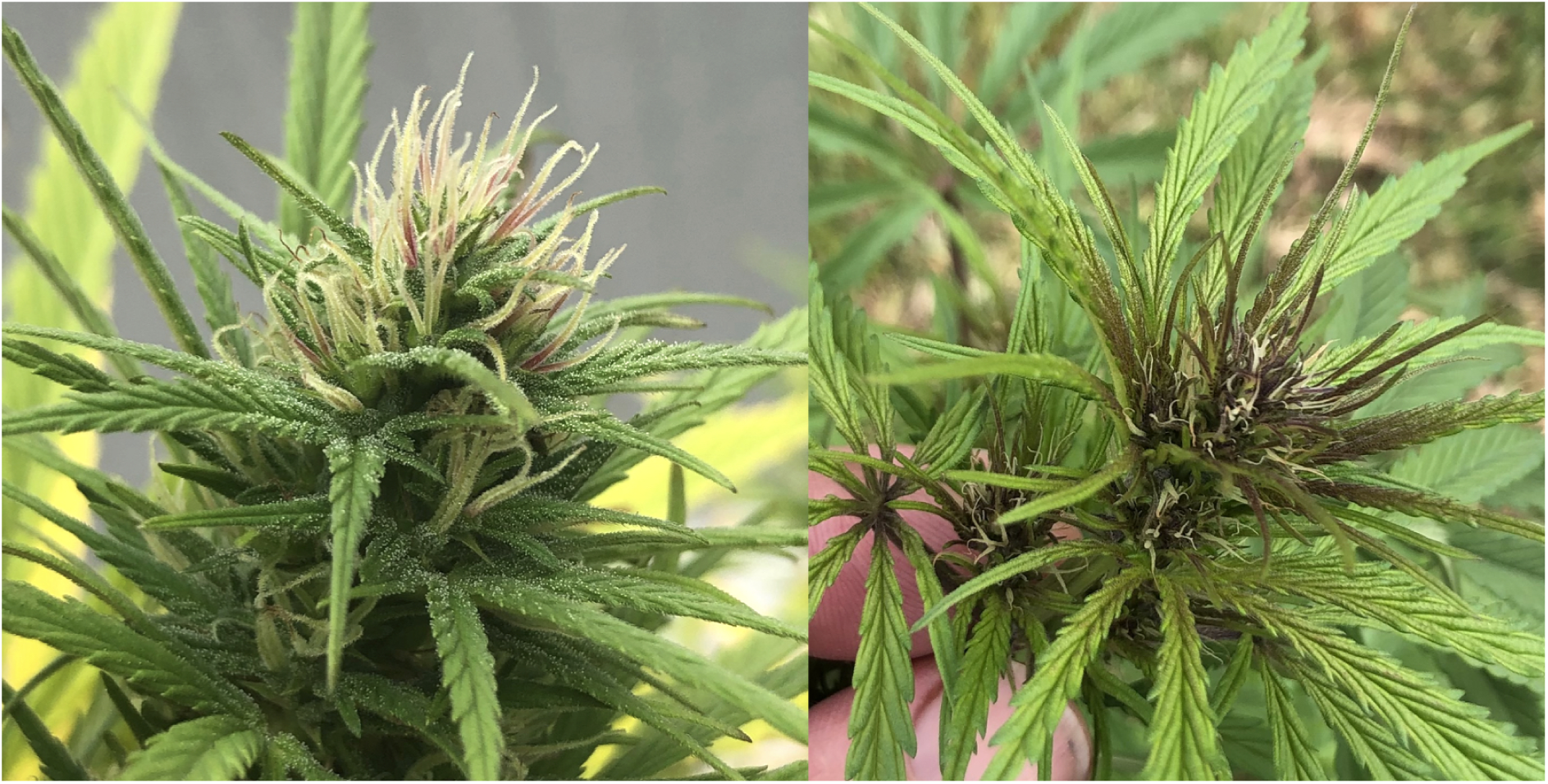
The Punto Rojo phenotype may describe anthocyanin deposition in the calyxes (left) or new shoots (right). Photos by author.

Cherry Pie (Fig. 2) is one of several Type III (CBD-dominant) strains in the Cherry family, bred in the American state of Colorado following legalization. Cherry Blossom^11^ and Cherry Wine^12^ have been the subjects of recent reports, and the NCBI reference for Cannabis, CBDRx^9^, falls into this clade as well. All display fast flowering and high CBD content, as well as a pleasant cherry aroma. The CP-16 individual was selected for consistently containing less than 1% THC at maturity, which enables its registration as non-psychoactive under Colombian law. This permits unlimited cultivation for any licensed cultivator, without diminishing Colombia’s share of the global THC quota established by the United Nations Office on Drugs and Crime.

**Figure 2.**
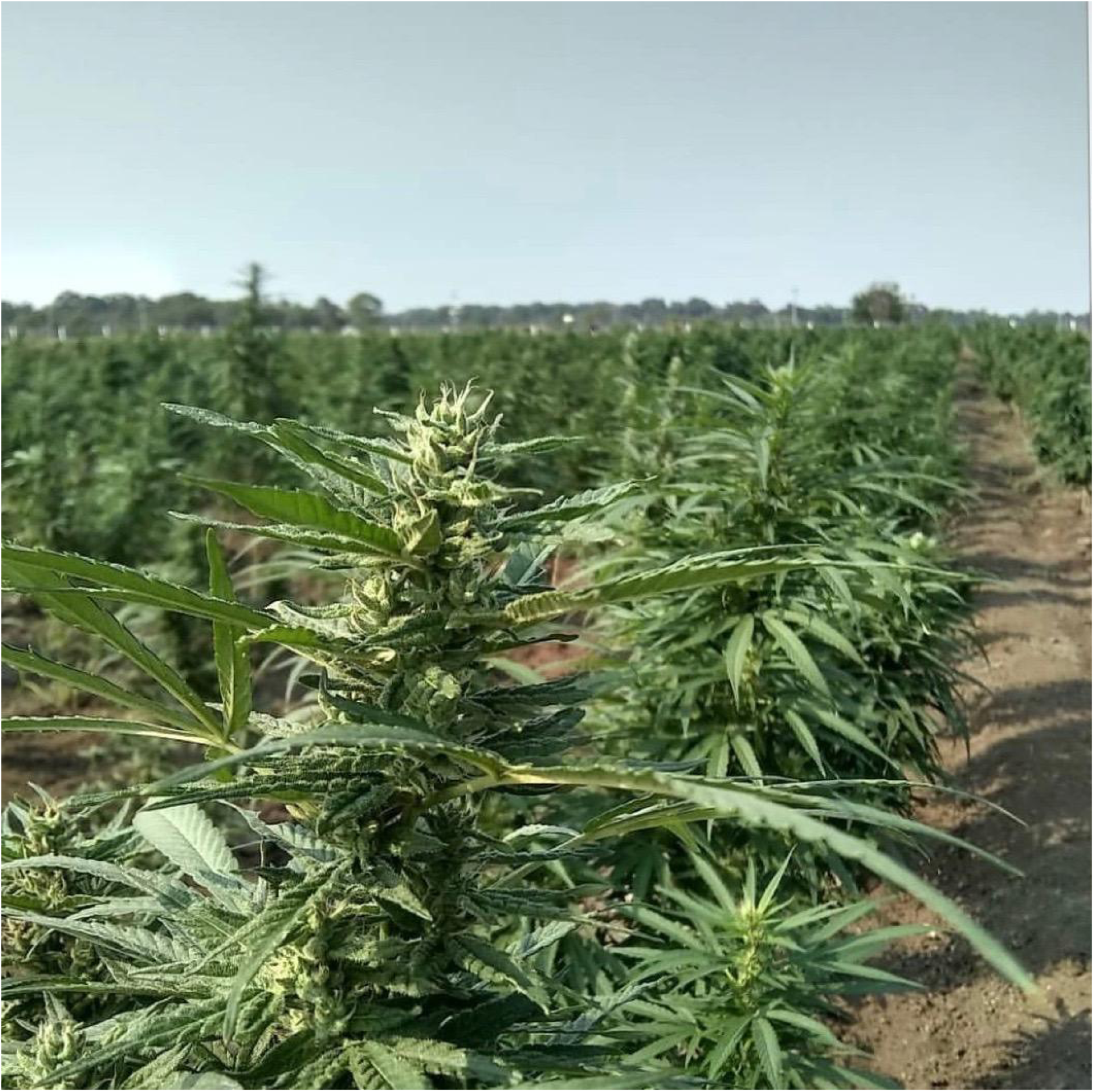
Clones of Cherry Pie #16 flowering in Fuente de Oro, Meta, Colombia. Photo courtesy of Medcann Pharma.

To facilitate comparative genomics, and establish a genome-wide resource for trait mapping and marker development, we assembled both parental haplotypes of this wide cross, via trio-binning of Oxford Nanopore reads. These fully-phased chromosome-scale assemblies have good contiguity and gene completeness, and can provide accurate catalogs of important gene families, specifically the disease resistance genes (R-genes) and terpene synthases (TPS).

### Biosynthesis of cannabinoids and terpenes

Cannabis produces two major cannabinoids (CN), cannabidiol (CBD) and tetrahydrocannabinol (THC). Minor cannabinoids, such as cannabichromene (CBC) and the propyl analogues cannabidivarin (CBDV) and tetrahydrocannabivarin (THCV), are also present at much lower levels. These metabolites are thought to modulate the behaviour of seed dispersers, and are primarily found in capitate stalked trichomes that cover the surface of female bracts and small leaves^13^. In wild cannabis, cannabinoids frequently constitute 0.3-15% of the mass in mature dry flower^14,15^, while in modern drug cultivars, reported CN content of 20-25% is common^16^. However, it is necessary to note that, in the newly legal market, technical inflation of CN content has been shown to be widespread^17^.

CBD and THC are synthesized in their acid form, from the common precursor cannabigerolic acid (CBGA), by the enzymes *CBD ACID SYNTHASE* (CBDAS) and *THC ACID SYNTHASE* (THCAS). The acid forms are water soluble and have some medical relevance^18,19^, but the neutral forms, which result from non-enzymatic decarboxylation, are more relevant to mammalian health and especially mood, because they can cross the blood-brain barrier. For clarity, here we use CBD and THC to refer to both the acid metabolite and its neutral degradation product.

The trait of major CN ratio is monogenic, with 95% of variability in CBD:THC ratio controlled by the B locus^20^. There exist two codominant alleles, B_D_ that encodes the gene for CBD, and B_T_ that encodes the gene for THC. Therefore, there are three genotypes, known as Types I, II, and III. Type I is B_T_/B_T_ and produces a CBD:THC ratio of about 1:30, Type II is B_T_/B_D_ and produces a ratio of about 6:5, with significant heritable variation in both directions, and Type III is B_D_/B_D_ and produces a ratio of about 23:1^15^. These ratios are strongly supported by *in vitro* expression in yeast^21^; however, *in vivo* the biological reality is less clear.

In recent years, long-read sequencing has revealed that these alleles span several megabases, contain one active enzyme and as many as 24 pseudogenes^22^, and do not recombine^9^. However, elucidating the precise structure has been difficult, because the degenerate paralogs are separated by tandem repeats of subunit length 30-45 kb^9^, which typically do not resolve accurately when assembled from reads of 15-20kb.

The total cannabinoid content shows continuous variation, and correlates with the type^13^ and size^23^ of trichomes produced. While trichome density has shown to correlate with essential oil content in hop^24^, at present no such correlation has been demonstrated in Cannabis^25^. Metabolically, the rate of CN accumulation is limited by production of the CBG precursor, which derives from a condensation of olivetolic acid with geranyl pyrophosphate^26^.

In addition to cannabinoids, Cannabis trichomes are also a rich source of terpenes, with flowers frequently having a terpene content of 2% dry mass^16^. These volatile aromatics are thought to modulate the psychological effects of CNs via unknown mechanisms^27^, as part of what is known as the ‘entourage effect’^28^. In animal models, inhalation of terpene-rich essential oils from lemongrass^29^, lavender^30^, and sweet orange^31^ has been shown to have anxiolytic and antidepressant effects, which may corroborate the experience of humans who inhale terpene-rich cannabis vapour. Specifically, recent research has shown that terpene content correlates strongly with retail labelling of Cannabis flowers^32^, which illustrates its importance to consumer preference.

A typical Cannabis sample will contain ten or more distinct terpenes^33^. This complex profile, or ‘terptype,’ is known to be polygenic^34^ and also environmentally labile^35^, due to their involvement in stress responses^36^, particularly plant-plant^37^ and plant-animal communication^38–40^. Several independent datasets support a model where the diversity in terptype can be divided into three ‘hyperclasses’: profiles dominated by myrcene, terpinolene, or limonene^41^. Because these profiles are heritable, medically relevant, and important to consumer preference, the incorporation of molecular markers into a terpene breeding program could be of great value.

Similarly to the R-genes described below, terpene synthases (TPS) are also highly diverse^42^, rapidly evolving^43^, and found in clusters of paralogs^44^.

### R-genes

The basis for plant pathology is the ‘gene-for-gene’ interaction^45^. In this model, monogenic resistance is conditioned by a single host gene that recognizes a specific protein (or other metabolite) produced by the pathogen. Broadly, transmembrane receptors detect apoplastic molecules common to classes of pathogens (pathogen associated molecular patterns, PAMPS) in pathogen-triggered immunity (PTI), while in effector-triggered immunity (ETI) effector molecules (proteins in all cases to date) delivered into the cell to modulate plant function are recognized by intracellular receptors that are highly specific. In filamentous pathogens effectors are typically delivered through haustoria, specialized structures that invade the apoplast and form extensive and very close associations with the cell membrane. Upon perception of the cognate effector, the R-protein can trigger a hypersensitive response (HR), in which programmed cell death (PCD) leads to a zone of dead, desiccated tissue around the point of infection that prevents further hyphal growth. Other strong defense mechanisms are also possible. PCD differs from necrosis or senescence^46^ in that the plant hastens its response by not making the effort to relocate expensive metabolites, and instead withdraws water as quickly as possible, so that even a necrotroph is foiled. This response has broad applicability, foiling various microbes, nematodes and other small herbivores, and by withholding water can even prevent the development of butterfly eggs laid on the leaf’s surface^47^.

Thus, predicting the R-gene repertoire of an assembled plant genome is of great importance to biotic stress research and for breeding more resistant crops. However, as these genes occur in dense clusters of paralogs^48,49^ they are difficult to map, and as they are the most diverse plant genes^50^ and evolve rapidly^51^, they are difficult to predict as well. Structurally, these R-genes contain a Toll/Interleukin-1 Receptor-like (TIR) or coiled-coil (CC) signalling domain, a nucleotide binding site (NBS) that in the absence of pathogen is ADP-bound and represses signalling activity, and a leucine rich repeat (LRR) that is capable of de-repressing signalling upon perception of the cognate ligand^52^. Upon perception the LRR induces a conformational change in the NBS that ejects ADP and enables ATP to bind, thereby deploying the signalling moiety. This class of protein are known as Niblrrs or NLRs (for Nucleotide binding, Leucine rich Repeat), with subtypes CNL and TNL respectively denoting a coiled-coil or TIR signalling domain. Around 100 of these genes have been cloned and verified as genuine R-genes, with the majority of that population coming to light since 2015.

Most NLR prediction pipelines employ a Hidden Markov Model (HMM), in which a collection of known sequences are used to derive a set of motifs that include both conserved and ambiguous positions^53^. This motif set is then used to decorate an unannotated genome or set of predicted proteins, which is then filtered for sequences containing the diagnostic motifs in the proper order and proximity^54,55^.

Among cultivated dicots, the number of R-genes may range from 47-1157^56^. As sequencing technology continues to become more accurate and less expensive, pan-genomic analyses will facilitate the identification of novel resistances^57,58^. A recent survey of the ‘pan-NLRome’^59^ characterizes the population of NLRs in Arabidopsis. By targeted sequencing of 65 accessions spanning the natural range of the species, and counting NLRs in each, the authors determine that choosing any 40 accessions tends to return very nearly the entire population of NLRs.

This area of Cannabis biology remains largely unstudied, with, to the best of our knowledge, only one R-gene having been mapped^60^. PM1 confers qualitative resistance to powdery mildew and lies within a cluster of about 10 NLRs at the distal end of chromosome 2. Introgression of such genes, especially from wild relatives, into elite breeding lines remains a major activity of plant breeders, and can greatly reduce the reliance on chemical controls.

In the legal market, most jurisdictions have stringent standards for permissible levels of agrichemical residues^61^. Particularly for concentrated cannabinoid products, recalls have been necessary to protect consumers from inhaling chemicals such as tebuconazole^62^ and piperonyl butoxide^63^. In the future, we hope that these and similar studies will enable the use of genetics, rather than chemistry, as an effective shield against pathogens, with concomitant reduction in the use of such compounds.

## Materials and Methods

### Breeding Materials

The sequenced individual was an F_1_ hybrid between the psychoactive Colombian landrace ‘Punto Rojo #3’ (PR) and the nonpsychoactive Coloradan line ‘Cherry Pie #16’ (CP). Both parental clones have been formally characterized and registered with the Instituto Colombiano Agropecuario (ICA) by Medicamentos de Cannabis SAS.

These parents contrastfor several important agronomic traits: height, flowering time, cannabinoid content, terpene content, and fungal susceptibility.

### Plant Growth

The F_0_s and the F_1_ were grown at the licensed farm of Medicamentos de Cannabis SAS near Fuente de Oro, Meta, Colombia, as approved by the Ministry of Justice in Resolution 1164 of 19 August 2021. At this latitude (3.47° N), the photoperiod is consistently 12h, and therefore always inductive for Cannabis flowering. At this altitude (400m), the average day and night temperatures are 30°C and 21°C. The F_0_ clones had previously been selected from seed and then propagated clonally. Clones were rooted in Oasis-type plugs under fluorescent lamps and then transplanted to 15L containers filled with 70% coco fibre and 30% worm castings, watered by hand, in a trailer about 2m x 4m, fitted with 2 1000w HPS lamps and an air conditioner set to 16C. The CP-16 female was induced to produce female (XX) pollen via two applications of 0.03% silver nitrate, at 0 and 7d of flowering, which was then blown towards a group of females, including PR, with the aid of an oscillating fan.

F_0_ clones were grown outdoors in 20L containers filled with a mix of 70% coco fibre and 30% worm castings, watered by hand as needed, and maintained in a vegetative state by providing supplemental illumination from 6pm until midnight.

F_1_ seeds were sown in two 144-cell trays and after 21d, 250 seedlings were transplanted to 3L containers filled with a mix of 70% coco fibre and 30% worm castings. These plants grew vegetatively for a total of 60d with 12h of sunlight and supplemental lighting from 6pm to midnight. They were next transplanted to the field at a density of 2 plants per square meter into holes amended with one handful of a mix consisting of 50% worm castings, 20% rock phosphate, 20% dolomite lime, and 10% Peruvian bat guano. The plants were rain-fed, with additional watering by hand as needed.

About 40d after transplant to the natural inductive photoperiod, an individual (PC-67) was chosen that was approximately average for the population in terms of height, flower development, leaf morphology, internode spacing, and degree of branching. As well, its flowers produced an aroma that evoked both the red fruit odor of Cherry Pie and the citric tanginess of Punto Rojo. PC-67 was cloned and propagated vegetatively, and about 12 weeks later, new shoots consisting primarily of unexpanded leaves were sampled for DNA sequencing.

### DNA purification

DNA from the F_0_s was extracted from new shoots dried over silica with a Quick-DNA Plant/Seed Miniprep Kit (Zymo Research, Irvine, California, USA).

For the F_1_, HMW DNA was purified from clean nuclei as described previously^64^ and then size-selected via the Short Read Eliminator XL kit (Circulomics, Baltimore, Maryland, USA). Several replicates were combined to yield sufficient quantity. DNA concentration and purity were estimated through the use of NanoDrop 1000 (Thermo Scientific), and two additional ethanol washes on SPRI beads were performed to meet sequencing standards.

### F_0_ Illumina Library Prep and Sequencing

The F_0_s were prepared as Illumina TruSeq libraries and sequenced as part of a NovaSeq PE150 lane. Illumina libraries were purified with BBDuk^65^ to remove adapter sequences and filtered by quality at 14 and by length at 21. In CLC Genomics Workbench, trimmed reads were filtered against Cannabis chloroplast and mitochondrial genomes and then assembled into contigs. These drafts were then used as BLAST queries against a custom database comprising the genomes of 7 fungi known or suspected to be present in the field: *Aspergillus fumigatus*, *Botrytis cinerea*, *Cercospora beticola*, *Fusarium oxysporum*, *Pseudocercospora fijiensis*, *Pseudocercospora musa*, and *Sclerotinia sclerotiorum*. Following these results, the unfiltered trimmed reads were then mapped with BBSplit^65^ to the genomes of *P. fijiensis* and *P. musae*, as well as the Carmagnola mitochondrion and Yunma-7 chloroplast, in order to reduce contaminants.

### F_1_ Oxford Nanopore Library Prep and Sequencing

The HMW sample was analyzed for length distribution via Agilent Femto-Pulse. Then, an Oxford Nanopore library was prepared (ligation kit LSK-0110) and sequenced in one PromethION R9.4.1 cell. After 24h, a nuclease flush was performed, and the library was then reloaded and sequenced for another 72h. Basecalling was performed by Guppy 5.1.12 in ‘super-accurate’ mode.

All library preparation and sequencing took place at The Genome Center at the University of California, Davis.

### Genome Assemblies

#### Genome size estimation

21-mers were counted in both sets of F_0_ short reads with jellyfish^66^ and histograms evaluated with findGSE^67^ in homozygous and heterozygous mode, with the latter using expected homozygous coverage of 18 (exp_hom=18). This process was repeated with the trio-binned, NECAT^68^-corrected F_1_ long reads.

#### Assembly

Trio binning was performed with scripts written for the purpose^69^. Briefly, 21-mers were counted with KMC^70^, unique parental 21-mers were derived by ‘find-unique-kmers,’ and 21-mers containing homopentamer repeats were deleted with a simple *grep* command. These lists were then used with ‘classify_by_kmers’ to sort long reads into PR, CP, and unknown bins.

Binned reads were assembled into contigs with NECAT, and the unbinned were ignored. Assembly included all reads longer than 3 kb with the default parameters and ‘polish contigs=false’. Contigs identified as mitochondrial by NCBI were removed. Assembly transpired on an AWS EC2 ‘m6gd.metal’ instance, with 64 ARM cores and 256 Gb RAM.

#### Polishing

Each haplotype’s binned raw reads were filtered for quality at 7 and aligned to their assembly with Minimap2^71^, with options ‘-aL -z 600,200 -x map-ont’. One round of polishing then took place with Racon^72^ with the ‘-u’ option.

Next, the appropriate F_0_ short reads were mapped to each haplotype with BWA MEM^73^ and polished with Clair3, twice. In the first round, Clair3 used the options ‘--haploid_precise--no_phasing_for_fa,’ which only generates well-supported 1/1 calls. In the second round, all variants were called: 0/1 calls were deleted, 1/1 calls were applied, and where possible the shorter allele in 1/2 calls was applied with the command ‘bcftools consensus -H SR’

Finally, each assembly was polished 4 times with its F_0_ kmers with ntEdit^74^, using default settings and kmer lengths of 40, 26, 40, and 26.

Polishing and other post-assembly processing took place on a 2012 Mac Pro 5,1 with 2 Xeon X5690 processors and 64 Gb RAM.

#### Scaffolding

Scaffolding was performed with ntJoin^75^ with options ‘nocut=True’ and ‘overlap=False,’ and a maximum gap of 100,000 bp. The substrate was derived from the Salk Institute’s recent release of many phased haplotypes^76^, which was subsetted to include 8 drug haplotypes assembled with benefit of Hi-C libraries. PR and CP contigs were first aligned to each haplotype, and alignments inspected visually with dotplotly^77^. For each chromosome, the homolog with the most diagonal alignment was chosen. Then, a small number of additional substitutions were made to reduce interchromosomal translocations.

The superscaffolds ultimately used for each genotype are listed in Table 1.

**Table 1.** Superscaffolds used for arranging PR and CP contigs to chromosome scale.

|  | PR | CP |
| --- | --- | --- |
| 1 | Golden Redwood B | White Widow A |
| 2 | White Widow A | Boax B |
| 3 | Sour Diesel B | Golden Redwood A |
| 4 | White Widow B | White Widow B |
| 5 | Boax A | White Widow A |
| 6 | White Widow A | Sour Diesel A |
| 7 | Boax A | Golden Redwood B |
| 8 | Golden Redwood A | Boax A |
| 9 | Sour Diesel B | Golden Redwood B |
| X | White Widow A | Boax A |

Finally, the chromosome-scale pseudomolecules were aligned to the cs10 reference genome^9^ and where necessary reverse complemented to maintain a consistent orientation. For PR, chromosomes 2, 3, 4, 8, and 9 were reversed, and for CP, 1, 6, 7, 8, 9 and X.

#### Evaluation

Assemblies were evaluated for contiguity with the BBTools^65^ script stats.sh, for completeness with compleasm^78^ using the BUSCO^79^ eudicots_odb10 5.4.6 database, and for correctness with yak^80^ via the QV module, after generating F_0_ kmer databases with yak’s count module and k=21. Accuracy of phasing was measured with yak’s trioeval module.

#### Organelles

F_0_ short reads identified as organellar were mapped to the Yunma-7 chloroplast and Carmagnola mitochondrion with the Geneious Prime 2023.0.4 mapper using default settings. The consensus sequences for each were generated and appended to each long-read assembly.

#### Diploid assembly

To test diploid-aware assembly methods, drafts were assembled in PECAT^81^ and Shasta^82^. PECAT used the configuration for Arabidopsis (cfg_arab_ont) with some modifications. Briefly, PECAT’s block size for correction and assembly, and Minimap2’s index and minibatch size, were raised to 40Gb to enable true all-versus-all alignment; for correction, minimum coverage was lowered to 2 for correcting and to 8 for calling SNPs for haplotypes; for assembly, the contig duplication rate was set to zero and only contigs over 4kb were outputted; and for phasing, minimum coverage was lowered to 16. The primary and alternate assemblies were then concatenated and reseparated with purge_haplotigs^83^.

Shasta used the Nanopore-Phased-May2022 configuration, and its output was further processed to resolve haplotypes: Assembly-Detailed.gfa and parental 31-mer databases generated with KMC^70^ were analyzed with GFAse^82^ to produce unphased, maternal, and paternal FASTAs.

To visualize haplotype resolution, parental unique hapmer databases were calculated in Meryl (with k=20) and drafts measured for their content in Merqury. These assemblies went unpolished and so QV is not reported. Contiguity and completeness were measured as above.

Diploid-aware analyses were performed on the ‘pyky’ node of the ZINE high-performance compute cluster at the Pontificia Universidad Javeriana, which includes 192 CPUs and 2 Tb of RAM.

### Annotation

#### Whole genome

Gene annotations were transferred from the cs10 reference to these drafts with Liftoff^84^, with options ‘-f features.txt -chroms chroms.txt -copies -sc 0.99,’ where features.txt includes all annotation types except ‘regions’, chroms.txt lists the most likely homolog for each pseudomolecule, based on a preliminary synteny analysis with SyRI^85^, and ‘-copies -sc 0.99’ seeks to find paralogs that have at least 99% exonic identity to the primary annotation.

#### Cannabinoid Synthases

Cannabinoid synthases were predicted *ab initio* in the assemblies listed in Table 2 by using the ‘Annotate From…’ function in Geneious Prime 2023.0.4^86^, using the full-length CDS for either THCAS from Skunk #1^87^ or the 6-3 allele of CBDAS^15^, a similarity threshold of 85%, and the ‘All matching annotations’ option. Gene clusters were then visualized in Geneious Prime.

**Table 2:** Genome size estimates in Mb, derived from F_0_ short and binned, corrected F_1_ long reads.

| readset | findGSE (hom) | error-excluded | findGSE (het) | error-excluded |
| --- | --- | --- | --- | --- |
| PR-ilmn | 857.661 | 823.229 | 857.661 | 819.607 |
| PR-ONT-bin-corr | 827.774 | 784.051 | fail | fail |
| CP-ilmn | 78.037 | 42.676 | 994.974 | 955.885 |
| CP-ONT-bin-corr | 794.321 | 751.697 | fail | fail |

#### Terpene Synthases

The cs10 annotations were filtered for the presence of the following descriptive terms: farnesene, geraniol, germacrene, humulene, limonene, linalool, myrcene, nerolidol, pinene, terpene, terpenoid, or terpinolene. The 47 annotations thus labelled were then transferred with Liftoff to both drafts, with stringency relaxed via ‘-copies -sc 0.50,’ to locate any additional paralogs that have similar structure and share at least 50% exonic identity.

To predict products, a custom BLAST database was built in Geneious Prime 2025.1.2 using the amino acid sequences of 33 TPS characterized via heterologous expression (as summarized by Booth et al^88^, Supp. Table S1). Predicted TPS were queried against this database with blastx, and in some cases multiply aligned with Clustal Omega.

#### Resistance Genes

The NBS_712 HMM^89^, which covers the highly conserved nucleotide binding site (NBS) region of NLRs and was initially derived from the Arabidopsis genome^54^, was queried with BLAST against the cs10 reference to create an initial list of candidates. These regions were extracted, aligned with ClustalW^90^, and used to create a Cannabis-specific NBS Hidden Markov Model (CsNBS HMM), via the hmmbuild and hmmemit modalities of the HMMER^91^ software package. The DNA consensus of the HMM was then BLASTed against the PR and CP drafts and hits, after merging overlaps, were taken as putative NLR loci.

As well, the NLR-Annotator^92^ was used to make a set of predictions, and the intersection of the two callsets was taken, so that full-length gene predictions from NLR-Annotator verified by CsNBS HMM hits are reported.

### Comparison

Drafts of PR and CP were each scaffolded to and then aligned against the collection of pseudomolecules listed in Table 1. Alignments were performed with Minimap2 with options ‘-cx asm5 --cs --eqx’ and visualized as a dotplot with dotplotly^77^. The resultant PAFs were analyzed with SyRI^85^ with default options, and visualized as a synteny map by plotting the SyRI calls with plotsr^93^. The PR and CP haplotypes were also compared to each other, and visualized in Circos^94^.

## Results

### HMW gDNA

Each prep of one gram of young shoots provided about 4 ug high-quality DNA, with 260/280 of 1.8 and 260/230 of 2.0. Analysis via Agilent Femto-Pulse showed that this method retains many fragments over 100kb, and the steep decline in fragments below ∼19kb suggests that the Short Read Eliminator XL kit did function as advertised (Fig. 3).

**Figure 3.**
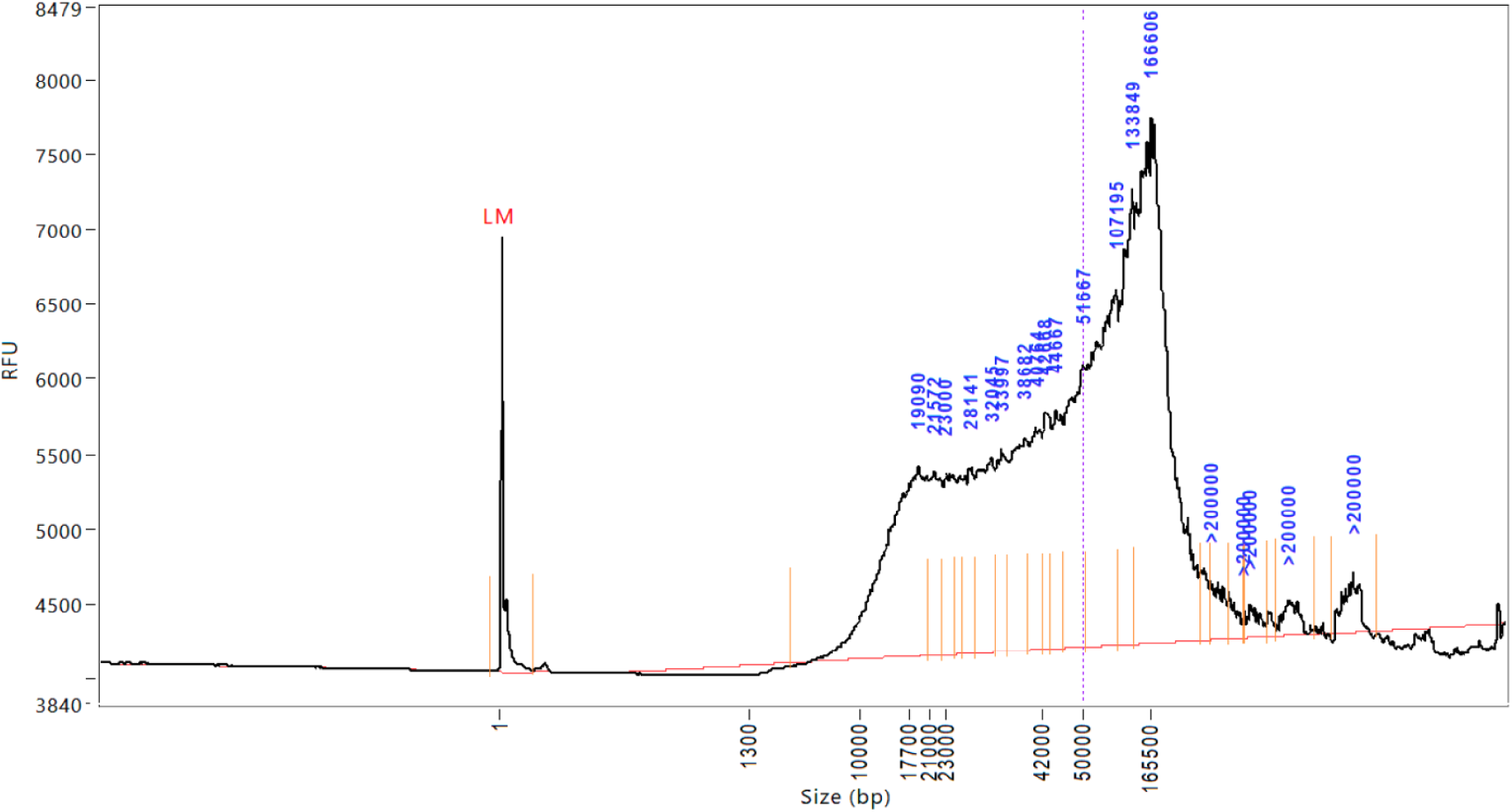
Agilent Femto-Pulse trace of gDNA fragment length distribution.

### F_0_ Illumina sequencing

The Illumina libraries yielded 52.6M and 47.6M read pairs for PR and CP.

Filtering the reads resulted in sets mapping to the chloroplast and mitochondria, as well as to two species of *Pseudocercospora*. Compared to a reference mitochondrion from the hemp line Carmagnola, PR contained 197 SNPs and CP 80. Compared to a reference chloroplast from Yunma-7, PR contained 9 SNPs and CP 125. About 0.5% of reads mapped to *Pseudocercospora*, with *P. musae* appearing to be about 50% more abundant than *P. fijiensis* in both F_0_s (data not shown).

After trimming and decontamination, 85.0% and 81.1% of base space remained, providing 16.7x and 14.4x of coverage for polishing.

### F_1_ Oxford Nanopore sequencing

The PromethION cell yielded 34.6 Gb of data, with an N_50_ of 23.6 kb, and 15.5% of bases contained in reads over 50kb.

### Estimation of Genome Size

Estimates of genome size were derived from both short and binned, corrected long reads. Results from findGSE are summarized in Table 2.

### Assembly

Assembly statistics, including the two drafts presented here, a recent NCBI upload, and three previously published chromosome-scale long-read assemblies^9,95,96^, are summarized in Table 3.

**Table 3:** Assembly statistics for PR, CP, and chromosome scale assemblies published since 2020. High marks are bolded where relevant. All BUSCO scores were newly calculated in compleasm with the eudicots_odb10 5.4.6 database.

| genotype | Punto Rojo | Cherry Pie | Abacus | Cannbio-2 | cs10 | JL |
| --- | --- | --- | --- | --- | --- | --- |
| citation | this study |  | - | Braich et al 2020 | Grassa et al 2021 | Gao et al 2020 |
| GenBank | JBDLLE000000000 | JBDLLD000000000 | GCA_025232715.1 | GCA_016165845.1 | GCA_900626175.2 | GCA_013030365.1 |
| long read tech | ONT |  | PacBio Sequel | PacBio SMRT | ONT | PacBio Sequel |
| LR coverage | 18x |  | 83x | 86x | 36x | **153x** |
| SR tech | pe150 |  | pe250 | - | pe150 | pe150 |
| SR coverage | 18x |  | **234x** | - | 100x | 118x |
| assembler | NECAT |  | unk. | HGAP4 | miniasm | Wtdbg, SMARTdenovo, Quickmerge |
| LR polish | Racon |  | unk. | HGAP4 | 3x Racon | blasr, Arrow |
| SR polish | 2x Clair3, 4x ntEdit |  | unk. | - | 3x Pilon | Pilon |
| scaffolder | ntJoin |  | Hi-C | RaGOO | Hi-C | Hi-C |
| scaffold reference | Salk assortment (Table 1) |  | de novo | cs10 | de novo | de novo |
| linkage map | - |  | - | - | Skunk x Carmen | - |
| haplotyping | trio-bin |  | unk. | - | - | - |
| # contigs | 867 | 1171 | 1023 | 8919 | **831** | 2978 |
| contig size (Mb) | 740 | 724 | 796 | 914 | 714 | 812 |
| contig N50 (Mb) | 2.12 | 1.65 | **3.17** | 0.17 | 2.14 | 0.51 |
| contig N90 (Kb) | 413 | 349 | 343 | 46 | **459** | 126 |
| **contig max (Mb)** | 9.84 | 7.86 | **16.96** | 1.56 | 10.06 | 2.87 |
| **# scaffolds** | 10 | 10 | 160 | 147 | **10** | 483 |
| **scaffold size (Mb)** | 794 | 774 | 797 | 914 | 854 | 813 |
| **scaffold N50 (Mb)** | 87.6 | 81.1 | 80.6 | 91.5 | 91.9 | 83.0 |
| **scaffold N90 (Mb)** | 66.4 | 68.9 | 63.0 | 71.6 | 64.6 | 69.1 |
| **N %** | 6.71% | 6.40% | 0.01% | 0.10% | 16.34% | 0.09% |
| **BUSCO** | **98.6%** | 94.5% | 97.2% | 97.1% | 94.0% | 92.8% |
| **single** | **2173** | 2149 | 2008 | 1373 | 2031 | 1794 |
| **duplicated** | 121 | **49** | 252 | 886 | 156 | 364 |
| **fragmented** | 7 | 21 | 9 | 16 | 16 | 23 |
| **missing** | **25** | 107 | 57 | 51 | 123 | 145 |
| **QV** | 29.2 | 28.8 |  |  |  |  |

#### Trio binning

The ‘classify_by_kmers’ script produced a PR bin containing 17,605 Mb of sequence in 1,238,187 reads, and a CP bin containing 15,942 Mb of sequence in 1,156,998 reads. The unknown bin contained 889 Mb in 322,224 reads, which did not assemble into contigs and were not analyzed further. The split among PR, CP, and unknown is 51.12%, 46.30%, and 2.58%.

After assembly and polishing, the switch rate and Hamming rate were estimated by yak trioeval as 2.9% and 2.0% for PR, and 3.0% and 2.2% for CP.

#### Contiguity

The drafts of Punto Rojo and Cherry Pie contain 1327 and 1881 contigs, with N_50_ of 2.0 and 1.6 Mb, contig N90 of 321 and 231 Kb, and a longest contig of 9.8 and 7.8 Mb.

Scaffolding PR and CP with ntJoin resulted in placement of 97.7% and 96.4% of contig sequence on the 10 chromosome-scale pseudomolecules, with N content of 6.6% and 6.2%.

#### Completeness

PR and CP have compleasm BUSCO scores of 98.6% and 94.5%, with duplication ratios of 5.2% and 2.1%. The full BUSCO output is summarized in Table 3 and Figure 4.

**Figure 4.**
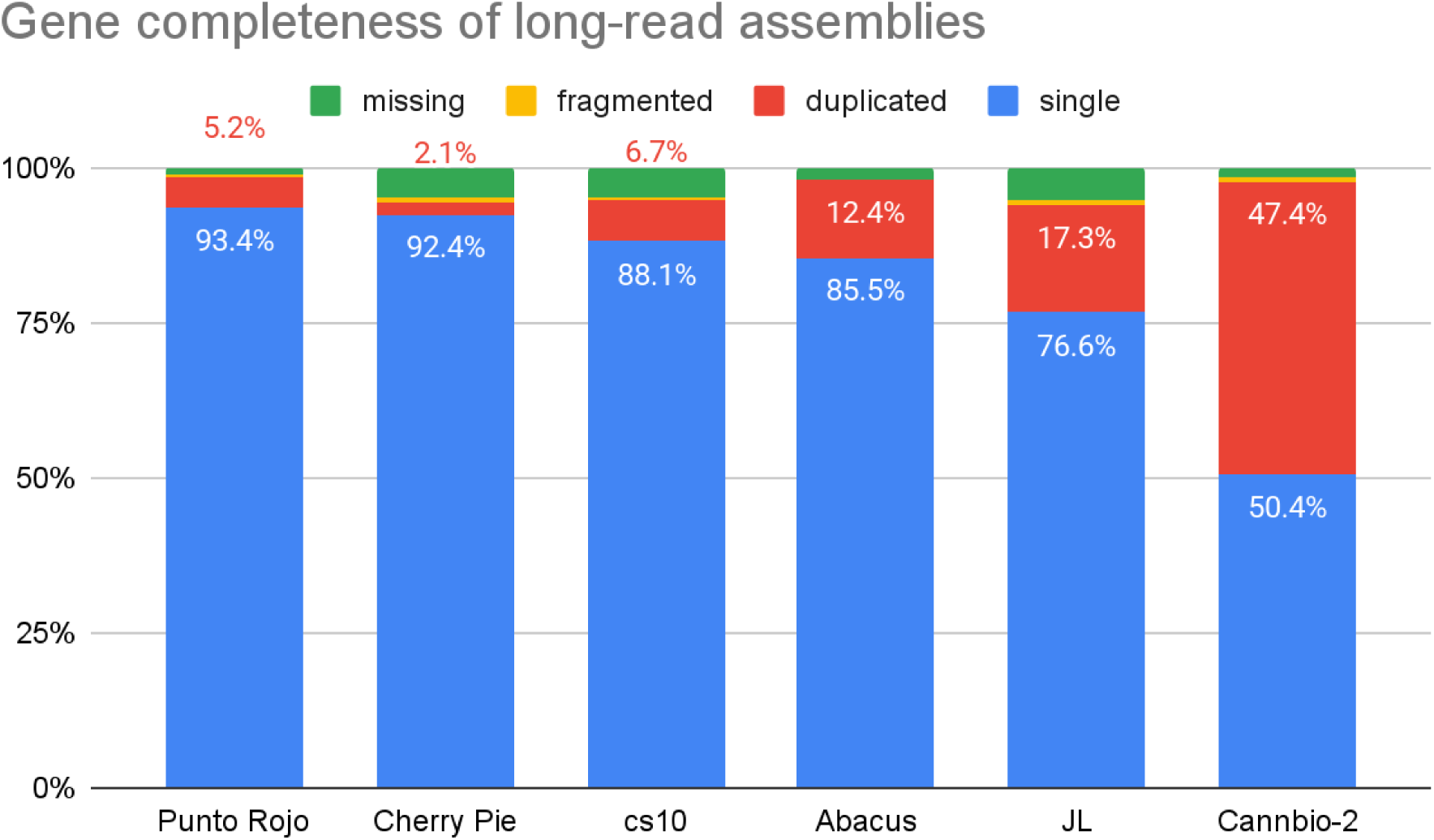
BUSCO scores of long-read Cannabis assemblies, with labels for single (blue) and duplicated (red) fractions. Previously published assemblies were newly evaluated with the eudicots_odb10 5.4.6 dataset.

#### Correctness

Yak eval estimates QV values for PR and CP of 29.2 and 28.8, corresponding in both cases to base level precision of 99.9%.

#### Diploid assembly

PECAT + purge_haplotigs produced a primary and an alternate assembly. Shasta produced a diploid draft which was subsequently binned by GFAse into maternal, paternal, and unknown compartments. The size, contiguity, completeness are reported in Table 4:

**Table 4:**
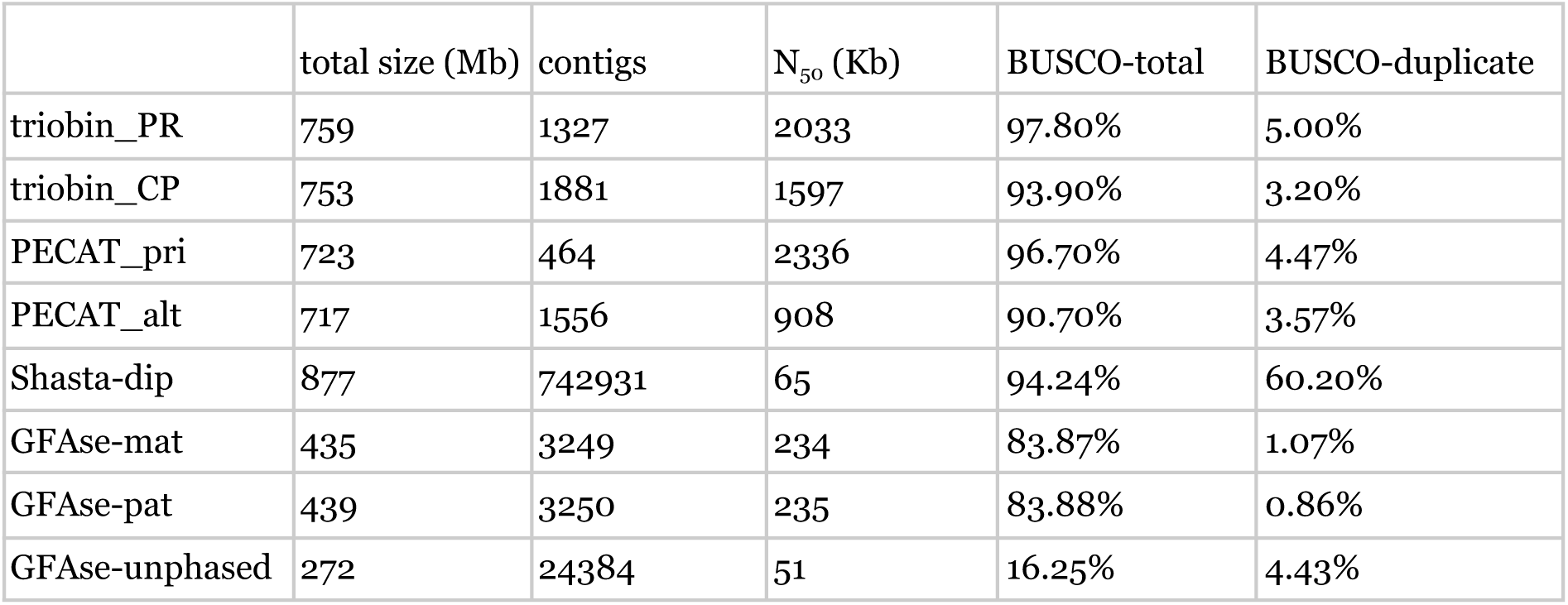
Assembly statistics for trio-binning, PECAT, Shasta, and Shasta + GFAse using F_0_ kmers.

| | total size (Mb) | contigs | $N_{50}$ (Kb) | BUSCO-total | BUSCO-duplicate |
| --- | --- | --- | --- | --- | --- |
| triobin_PR | 759 | 1327 | 2033 | 97.80% | 5.00% |
| triobin_CP | 753 | 1881 | 1597 | 93.90% | 3.20% |
| PECAT_pri | 723 | 464 | 2336 | 96.70% | 4.47% |
| PECAT_alt | 717 | 1556 | 908 | 90.70% | 3.57% |
| Shasta-dip | 877 | 742931 | 65 | 94.24% | 60.20% |
| GFase-mat | 435 | 3249 | 234 | 83.87% | 1.07% |
| GFase-pat | 439 | 3250 | 235 | 83.88% | 0.86% |
| GFase-unphased | 272 | 24384 | 51 | 16.25% | 4.43% |

Haplotype separation was visualized in Merqury, based on per-contig counts of parental short-read 20-mers as tabulated by Meryl (Fig. 5):

**Figure 5.**
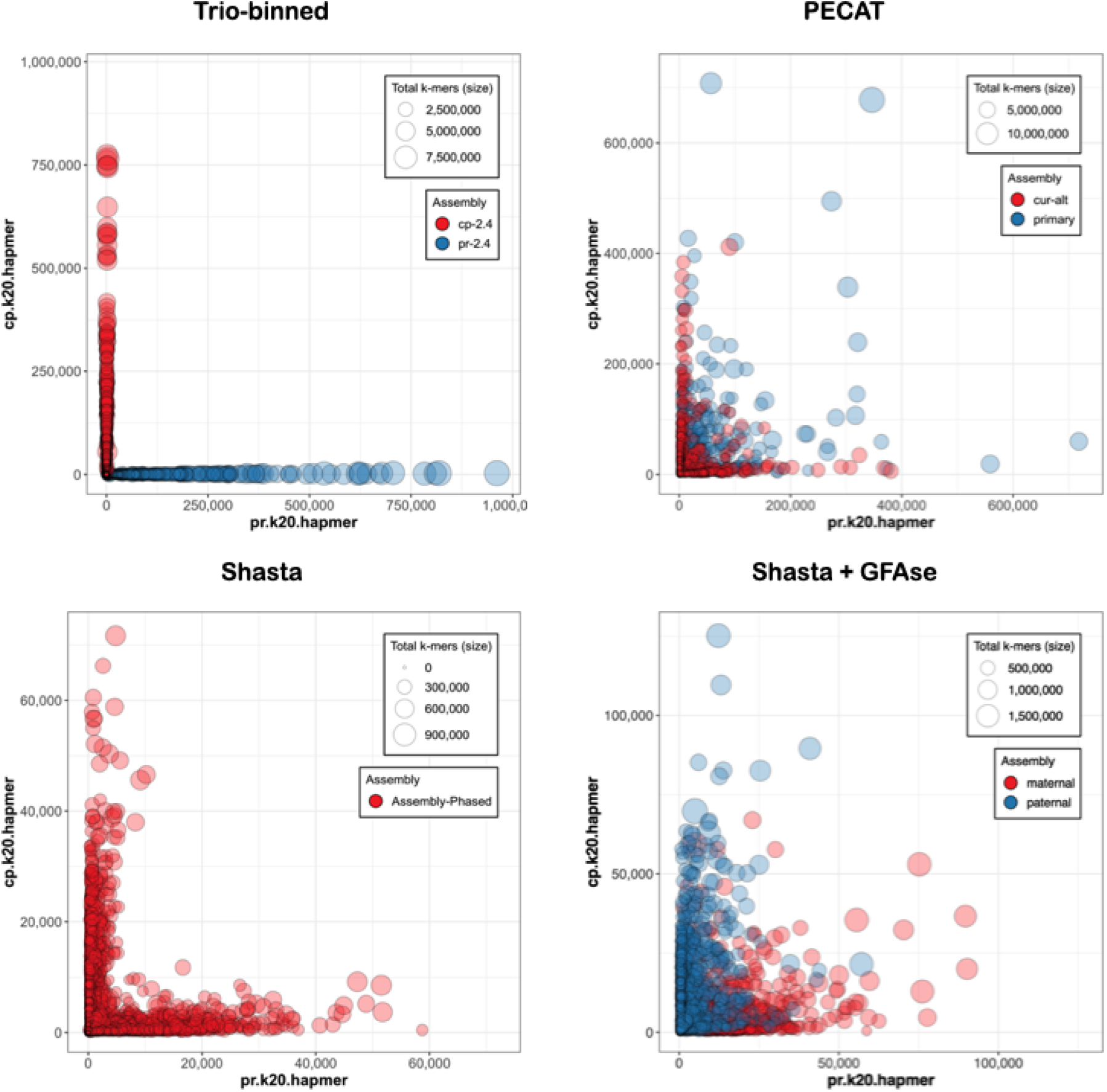
Merqury plots, where the X and Y axis represent the number of unique PR and CP 20-mers. Please note that scaling varies among drafts.

**Figure 6.**
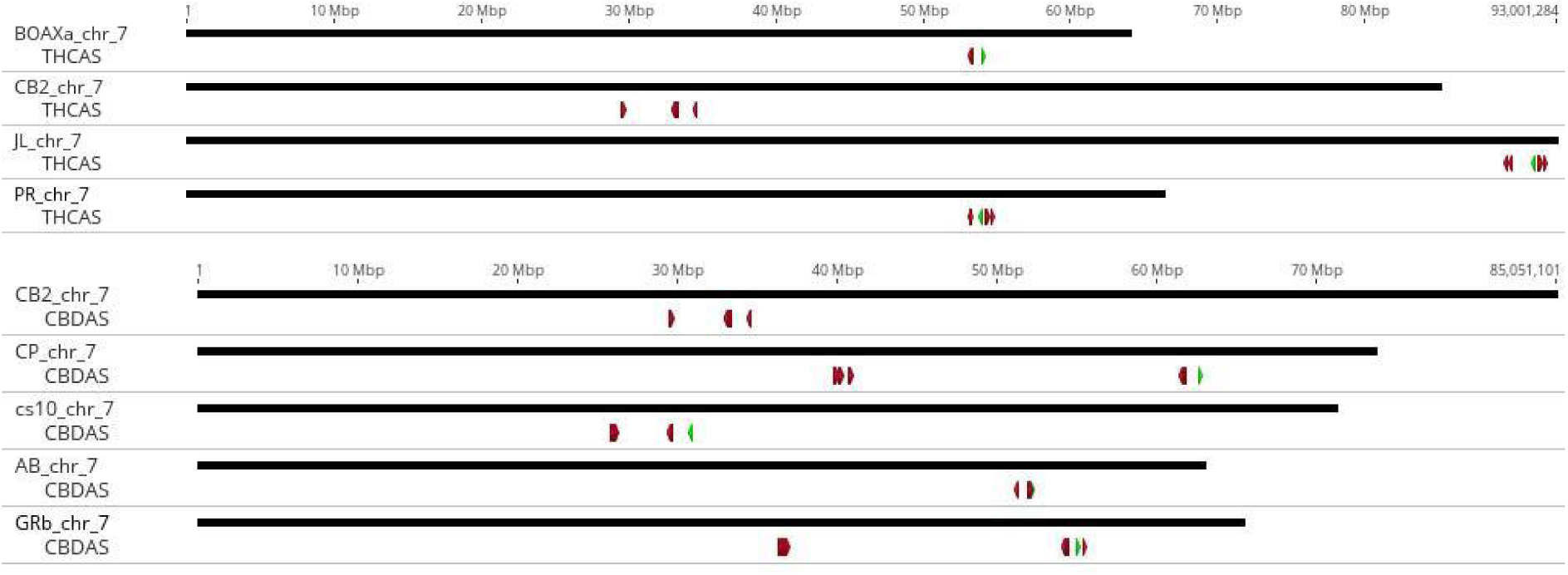
Visualization of chromosome 7 from the published assemblies and the haplotypes used for scaffolding (BOAXa for PR and GRb for CP). The active synthase is marked in green, while inactive paralogs are in red.

### Annotation

#### Liftoff

Nearly all of the reference annotations were able to be placed on both drafts. Table 3 summarizes the drafts’ annotations:

**Table 4a.**
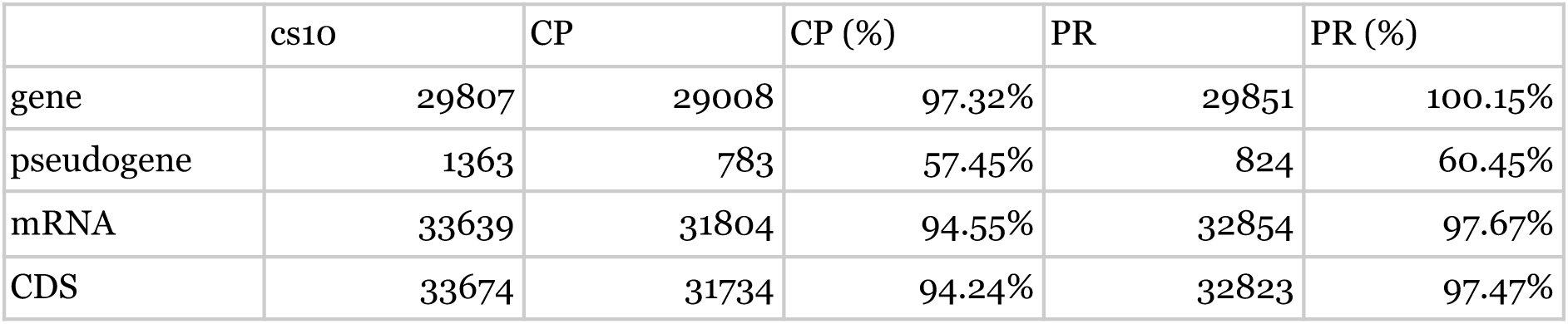
Accounting of cs10 annotations transferred to PR and CP with Liftoff, with option ‘-copies 0.99’.

#### CN synthases

The primary location for CN synthases, which includes 6-13 paralogs with identity from 85.3-99.9%, is the previously identified B locus^9,20^ on chr7, which varies in size, location, and copy number among assemblies (Table 5 and Figure 5). We note here that JL numbers its chromosomes in order of length, so that its chr1 is the homolog of chr7 in cs10 and the other listed assemblies. Because PR and JL do not include a CBDAS above 95% identity, and CP and Abacus do not include a THCAS above 95% identity, we report only the relevant CN synthase query and homology scores for paralogs of the putative active gene, which in all cases shares >99% identity with the query. However, we note that in no case does a query with the other CN synthase return a different copy number (data not shown). Because Cannbio-2 is a pseudohaploid representation of a B_D_/B_T_ genotype, its results are reported for both queries.

**Table 5.** Copy number variation among THCAS and CBDAS paralogs over 85% identity. Bolded cells indicate putative active synthases.

| genotype | PR | JL | CB2 | CP | Abacus | cs10 | CB2 |
| --- | --- | --- | --- | --- | --- | --- | --- |
| query | THCAS | THCAS | THCAS | CBDAS | CBDAS | CBDAS | CBDAS |
| coord (Mb) | 53.1-54.6 | 89.7-92.2 | 29.5-34.6 | 39.8-62.7 | 51.3-52.2 | 25.8-31.0 | 29.5-34.6 |
| 1 | **99.94%** | **99.21%** | **99.94%** | **99.94%** | **99.94%** | **99.94%** | 93.53% |
| 2 | 91.82% | 91.70% | 96.03% | 89.99% | 90.17% | 90.11% | 92.68% |
| 3 | 91.76% | 91.70% | 95.97% | 89.68% | 90.05% | 89.07% | 90.05% |
| 4 | 91.21% | 88.53% | 92.37% | 89.27% | 90.05% | 89.01% | 90.05% |
| 5 | 88.53% | 87.61% | 92.37% | 89.19% | 89.87% | 88.78% | 89.99% |
| 6 | 87.61% | 87.22% | 92.25% | 89.19% | 89.38% | 88.52% | 89.68% |
| 7 |  |  | 91.94% | 88.90% |  | 88.41% | 89.62% |
| 8 |  |  | 91.88% | 88.66% |  | 88.10% | 89.56% |
| 9 |  |  | 91.82% | 88.54% |  | 87.64% | 89.38% |
| 10 |  |  | 87.42% | 88.31% |  | 85.98% | 89.19% |
| 11 |  |  | 86.70% | 88.30% |  | 85.29% | 89.13% |
| 12 |  |  |  | 88.27% |  |  |  |
| 13 |  |  |  | 88.01% |  |  |  |

The arrangement of CBDAS copies appears to offer more variability. While most drafts contain all synthase copies in one cluster of 5Mb or less, CP has two clusters, both on chr7: a group of 5 containing the active synthase at 61.7-62.7 Mb, and a group of 8 paralogs with 88-89% identity that spans from 39.8 to 40.9 Mb. The Golden Redwood B haplotype to which it is scaffolded appears similar, but contains 7 and 10 copies in similarly situated clusters.

#### TPS

The annotations transferred from cs10 were mined for descriptions that included the name of a terpene. Table 6 summarizes those results:

**Table 6.** Counts of terpene synthases in cs10, PR, and CP.

|  | cs10 | PR | CP |
| --- | --- | --- | --- |
| farnesene | 1 | 1 | 1 |
| geraniol | 7 | 7 | 7 |
| germacrene | 8 | 7 | 6 |
| humulene | 6 | 6 | 6 |
| limonene | 2 | 2 | 2 |
| linalool | 4 | 4 | 2 |
| myrcene | 6 | 6 | 6 |
| nerolidol | 2 | 1 | 1 |
| pinene | 0 | 0 | 0 |
| terpene | 10 | 10 | 9 |
| terpenoid | 1 | 1 | 1 |
| terpinolene | 0 | 0 | 0 |
| TOTAL | 47 | 45 | 41 |

The TPS are unevenly distributed, with clusters of monoterpene and diterpene synthases lying in subtelomeric regions of chromosomes 5 and 6, respectively, as well as a cluster of probable monoterpene clusters on chromosome 9. We denote these as Major Terpene Clusters (MTC, Table 7), defined here as a group of at least 4 TPS genes separated from one another by no more than 2 Mb.

**Table 7.** Terpene synthases found in clusters. The TPS10 triplet in MTC5 includes one TPS10 and two TPS10-like predictions in both haplotypes.

| genotype | chr | start | stop | TPS | content |
| --- | --- | --- | --- | --- | --- |
| PR | 5 | 0.9 Mb | 2.6 Mb | 10 | 3x TPS10, 3x Myrcene, 2x Limonene, 2x Myrcene |
| CP | 5 | 1.4 Mb | 2.7 Mb | 10 | 3x TPS10, 3x Myrcene, 2x Limonene, 2x Myrcene |
| PR | 6 | 80.4 Mb | 82.9 Mb | 9 | 2x Humulene, 4x Germacrene, 3x Humulene |
| CP | 6 | 75.3 Mb | 78.3 Mb | 10 | 2x Humulene, 5x Germacrene, 3x Humulene |
| PR | 9 | 59.2 Mb | 59.4 Mb | 5 | 5x probable monoterpene synthase |
| CP | 9 | 62.9 Mb | 63.0 Mb | 4 | 4x probable monoterpene synthase |

To corroborate the predicted products, we queried a custom BLAST db, composed of 33 TPS characterized by heterologous expression, with the CDS of TPS found in cs10, PR, and CP. Where a gene contains multiple isoforms we took isoform X1. To quantify similarity we report the ‘Grade’, a proprietary metric within Geneious Prime that incorporates the length, e-value, and percent identity of the hit.

**Table 8.**
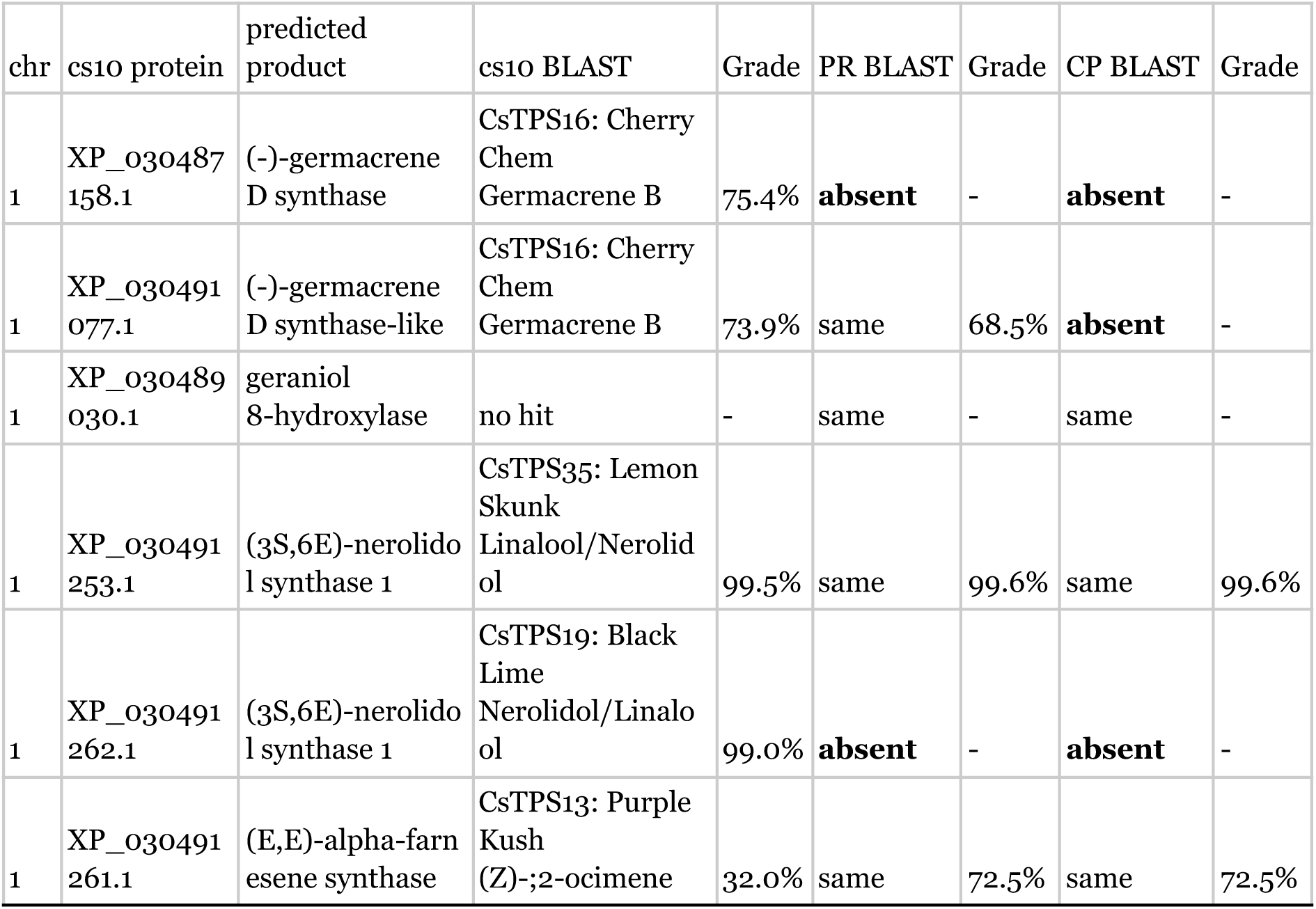
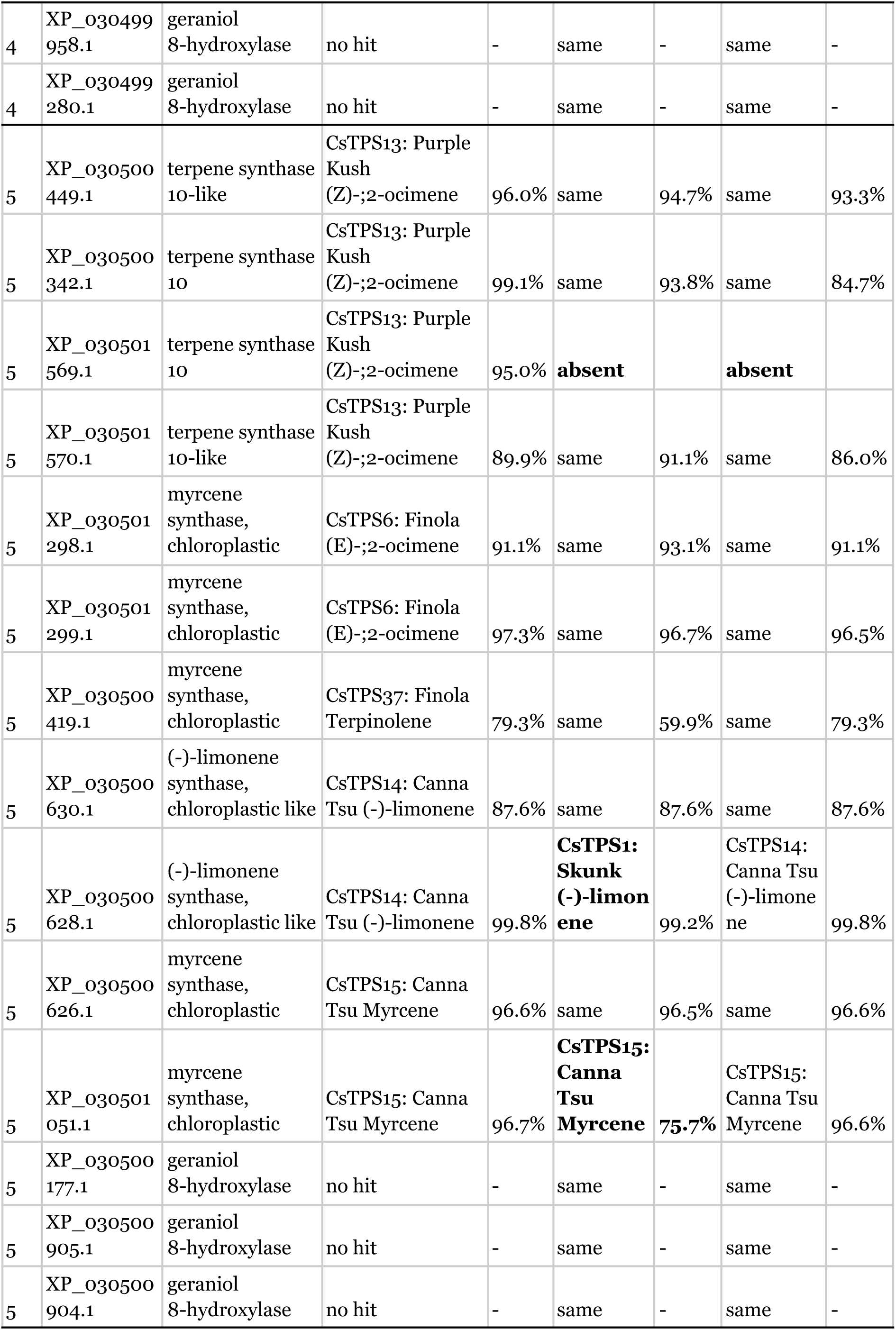
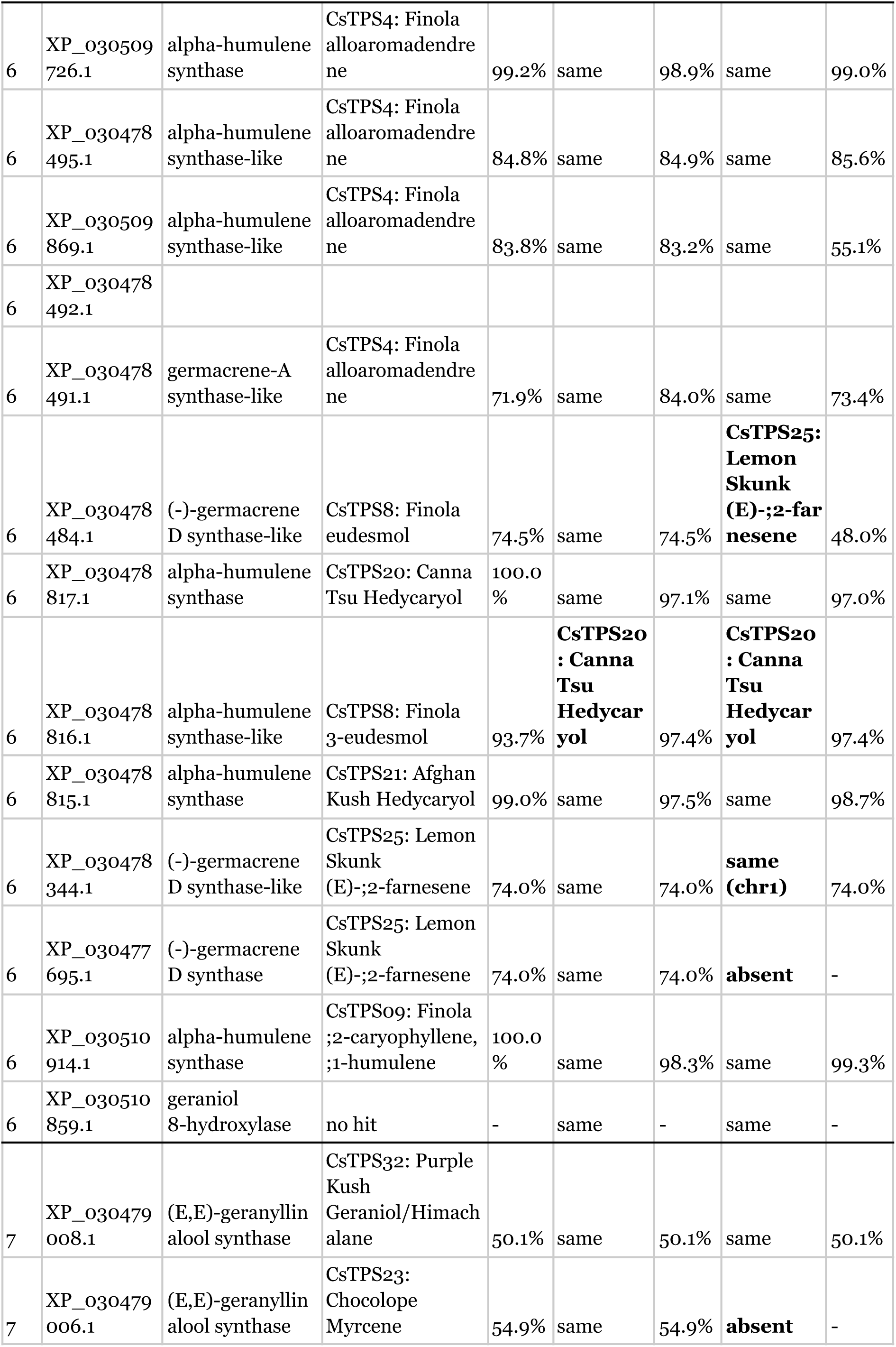
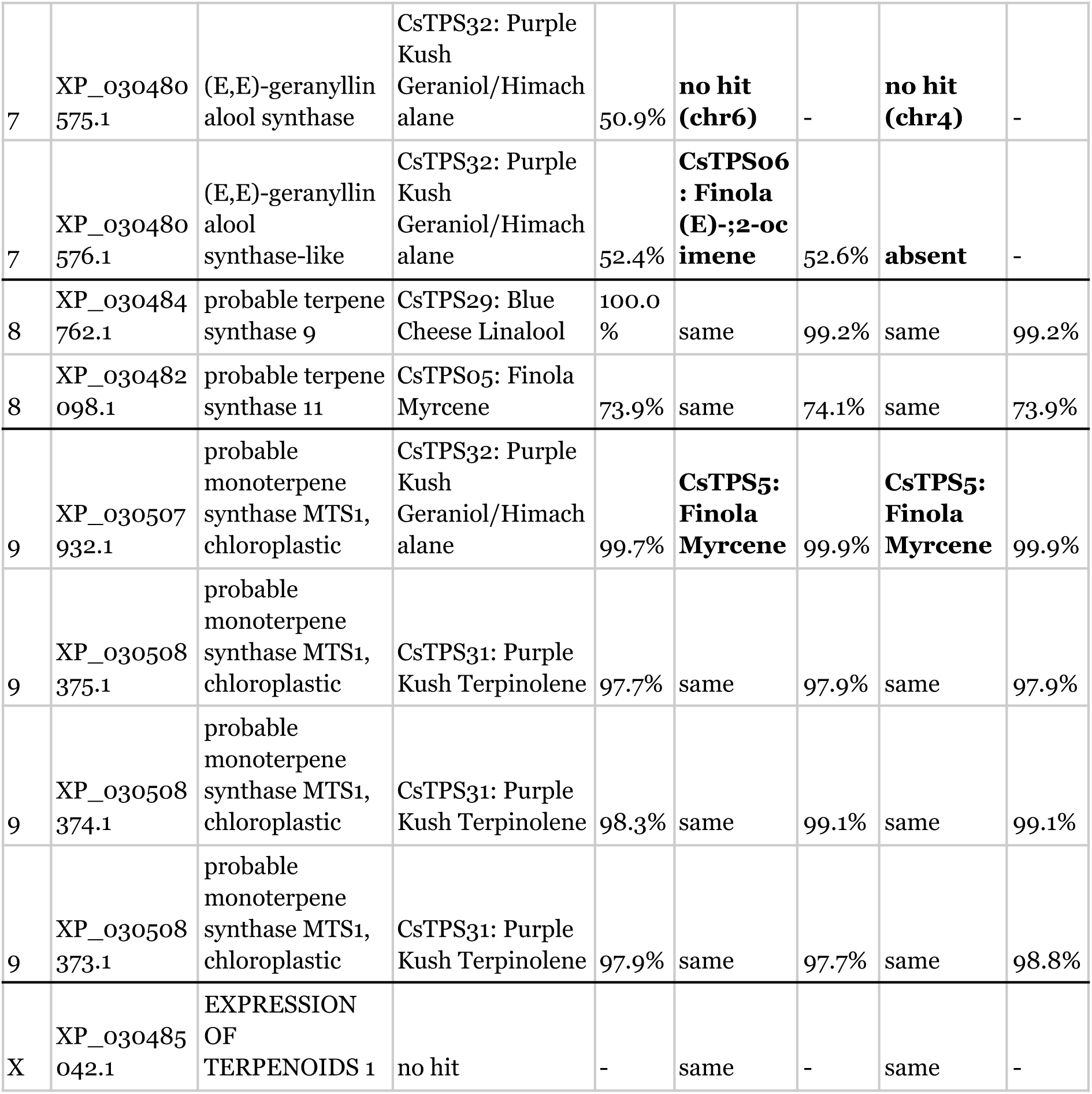
Best BLASTx hits of predicted TPS against 33 enzymes characterized via expression *in vitro*. Noteworthy variations bolded.

Aligning limonene synthases revealed a proline-serine transversion (Fig. 7) and aligning myrcene synthases revealed several nonsense mutations (Fig. 8).

**Figure 7.**
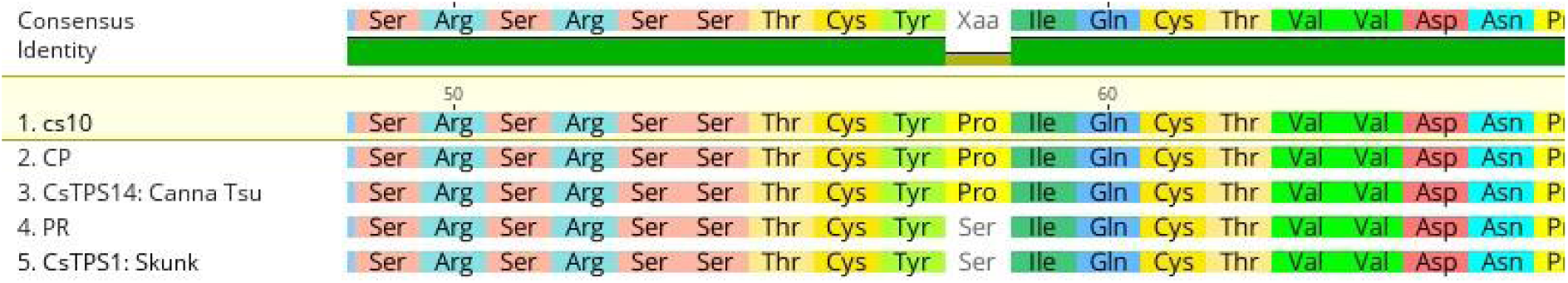
Clustal Omega alignment of limonene synthases from cs10, PR, CP, Canna Tsu, and Skunk.

**Figure 8.**
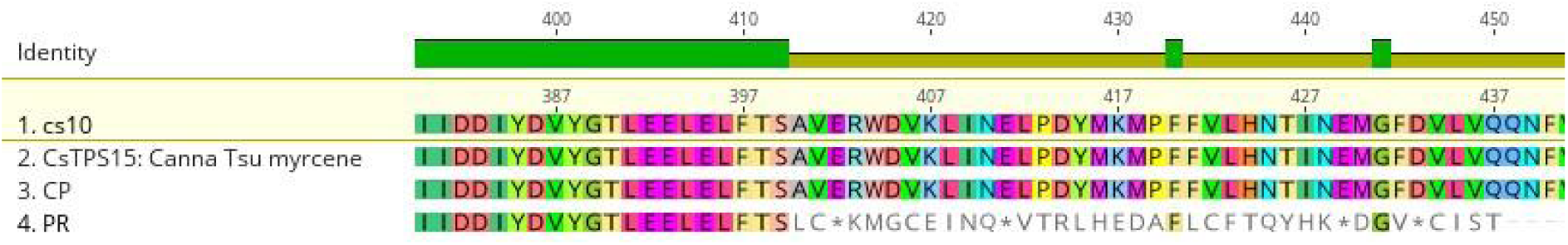
Clustal Omega alignment of myrcene synthases from cs10, PR, CP and Canna Tsu.

#### NLRs

We report 227 results in PR and 240 in CP, all of which are placed on the 10 chromosomes. Many of these predictions occur in clusters, which we call Major Resistance Clusters (MRC)^97^. Due to their more abundant and diffuse nature, we forego a formal definition and instead rely on a simple visual inspection. Typically, clusters have 5 or more members and an NLR density of at least one NLR per 2 Mb.

In PR, 176 NLRs are found in 9 clusters, and in CP, 188 in 11 clusters, representing 77.5% and 78.3% of the total (Table 9.) While most MRC have similar location and copy number between drafts, MRC1a has 8 NLRs in PR compared to just 2 in CP, and MRC5 and MRC7, which contain 4 and 14 NLRs in CP, appear to be absent from PR.

**Table 9.**
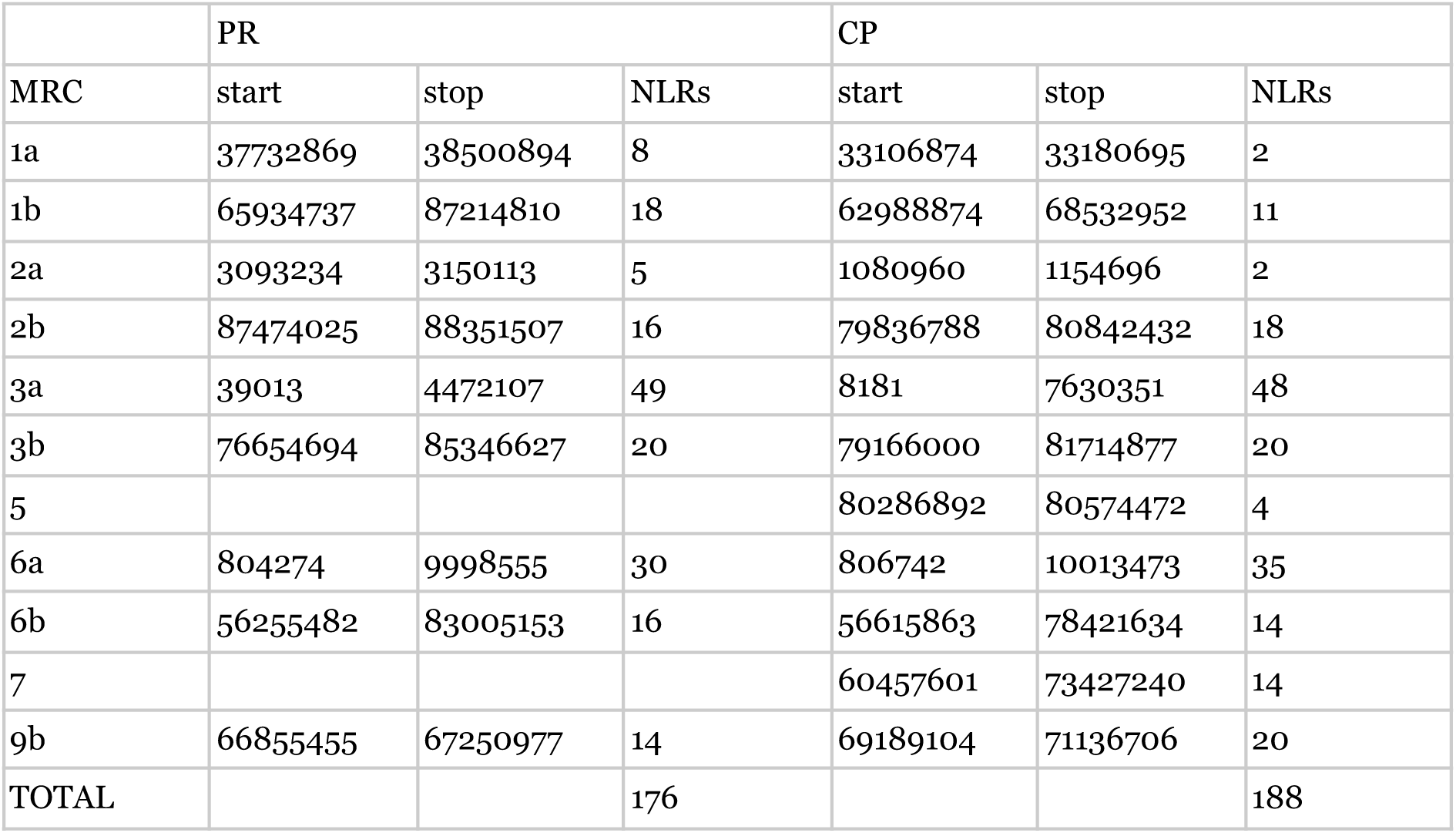
Location and copy number of major resistance gene clusters.

### Comparative Genomics

PR and CP were scaffolded to and then aligned against the set of chromosome-scale pseudomolecules shown in Table 1. SNPs and larger variants are summarized in Table 10.

**Table 10.**
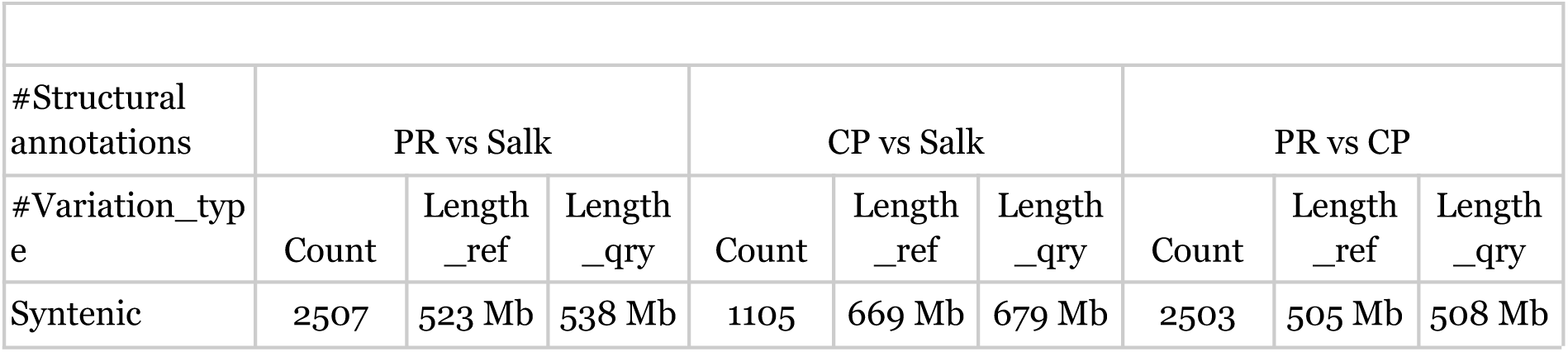
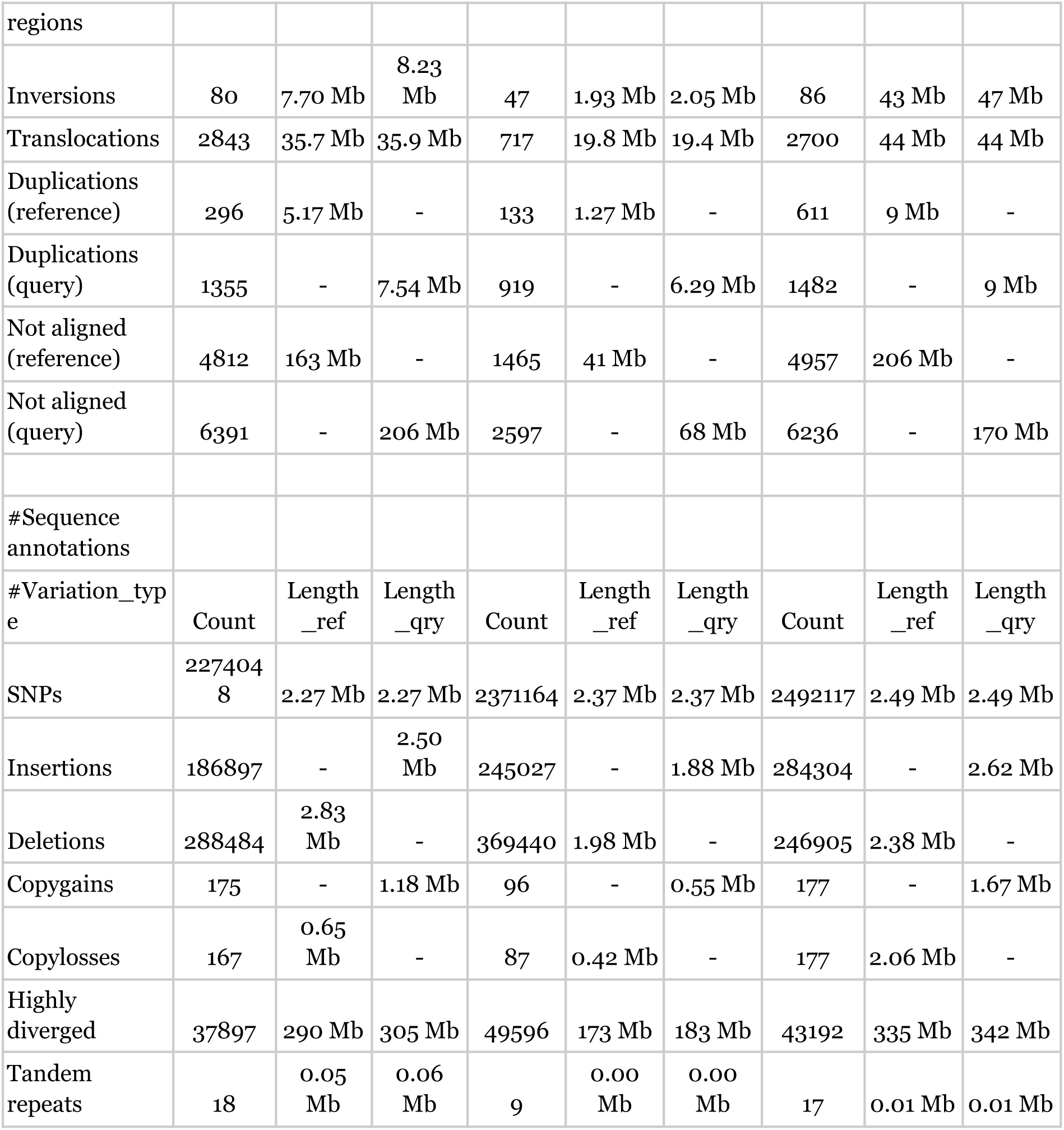
Structural and sequence variation as reported by SyRI for PR and CP.

To visualize collinearity, dotplots were generated for each draft relative to its scaffolding substrate (Fig. 9). As well, synteny between the two haplotypes was plotted with SyRI and plotsr (Fig. 10) and a Circos plot was generated that, in addition to synteny and interchromosomal translocations, includes tracks for contig boundaries, gene density, and the location of TPS and NLR genes (Fig. 11).

**Figure 9.**
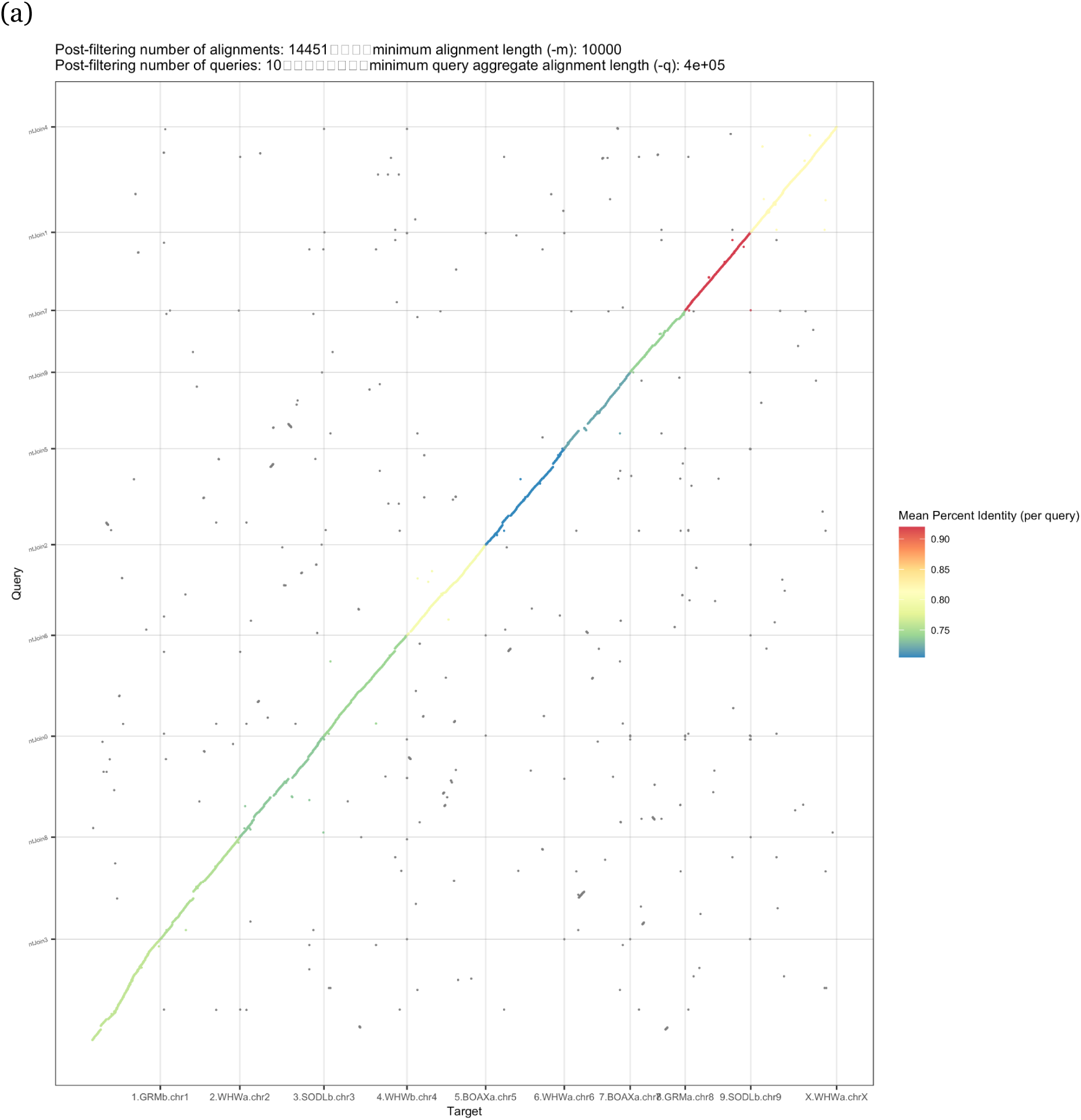
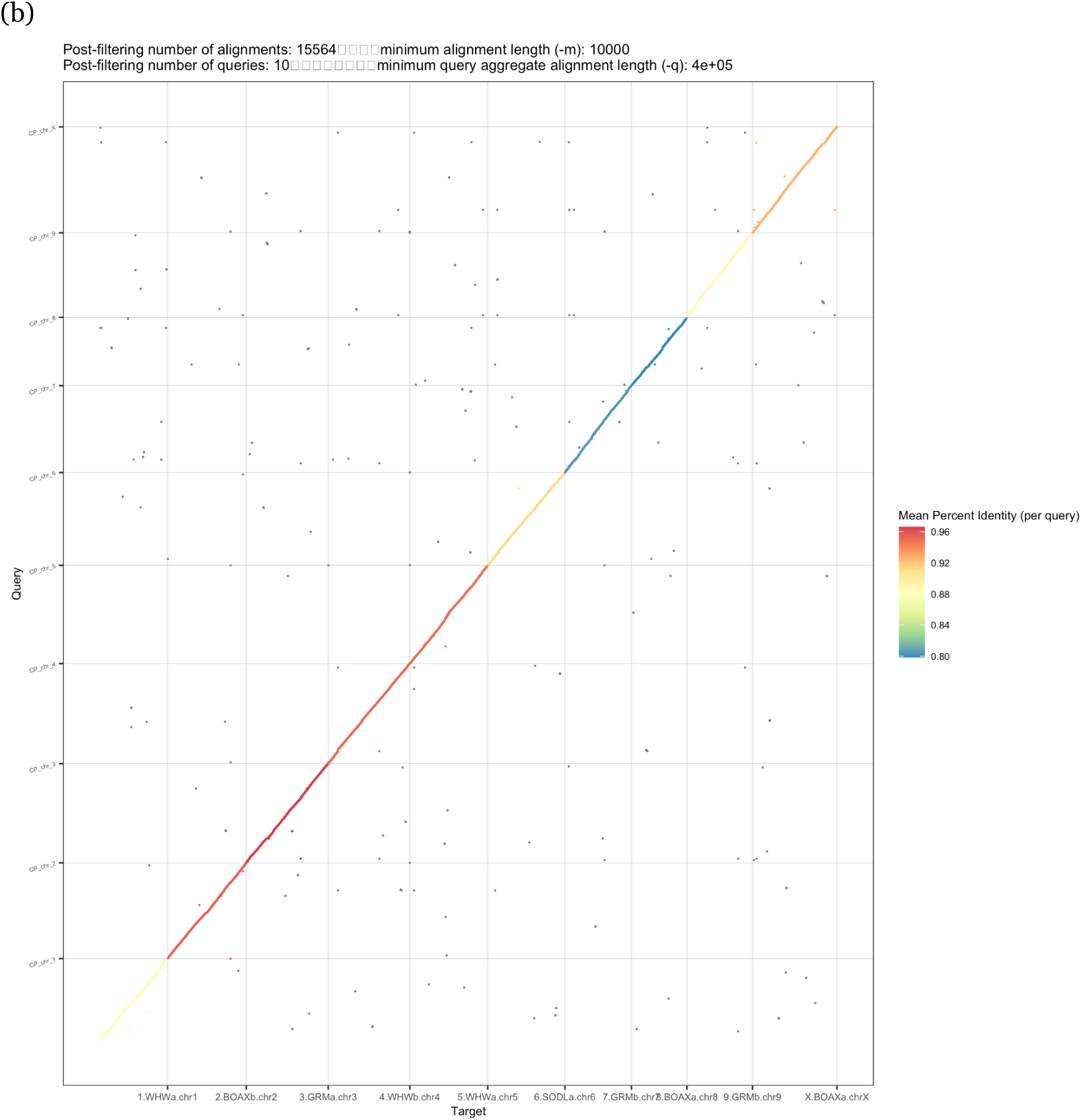
Dotplot showing alignment between the (a) PR and (b) CP haplotypes and their respective scaffolding substrates.

**Figure 10.**
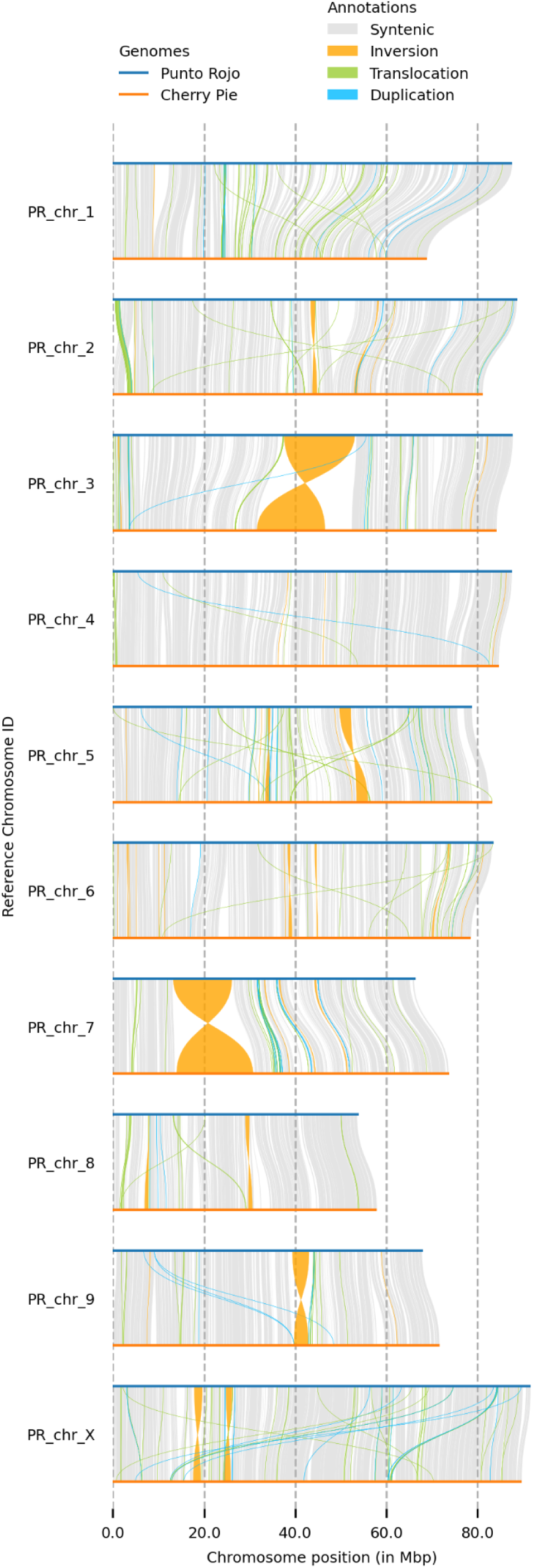
Synteny and rearrangement between PR and CP homologs, filtered above 100kb.

**Figure 11.**
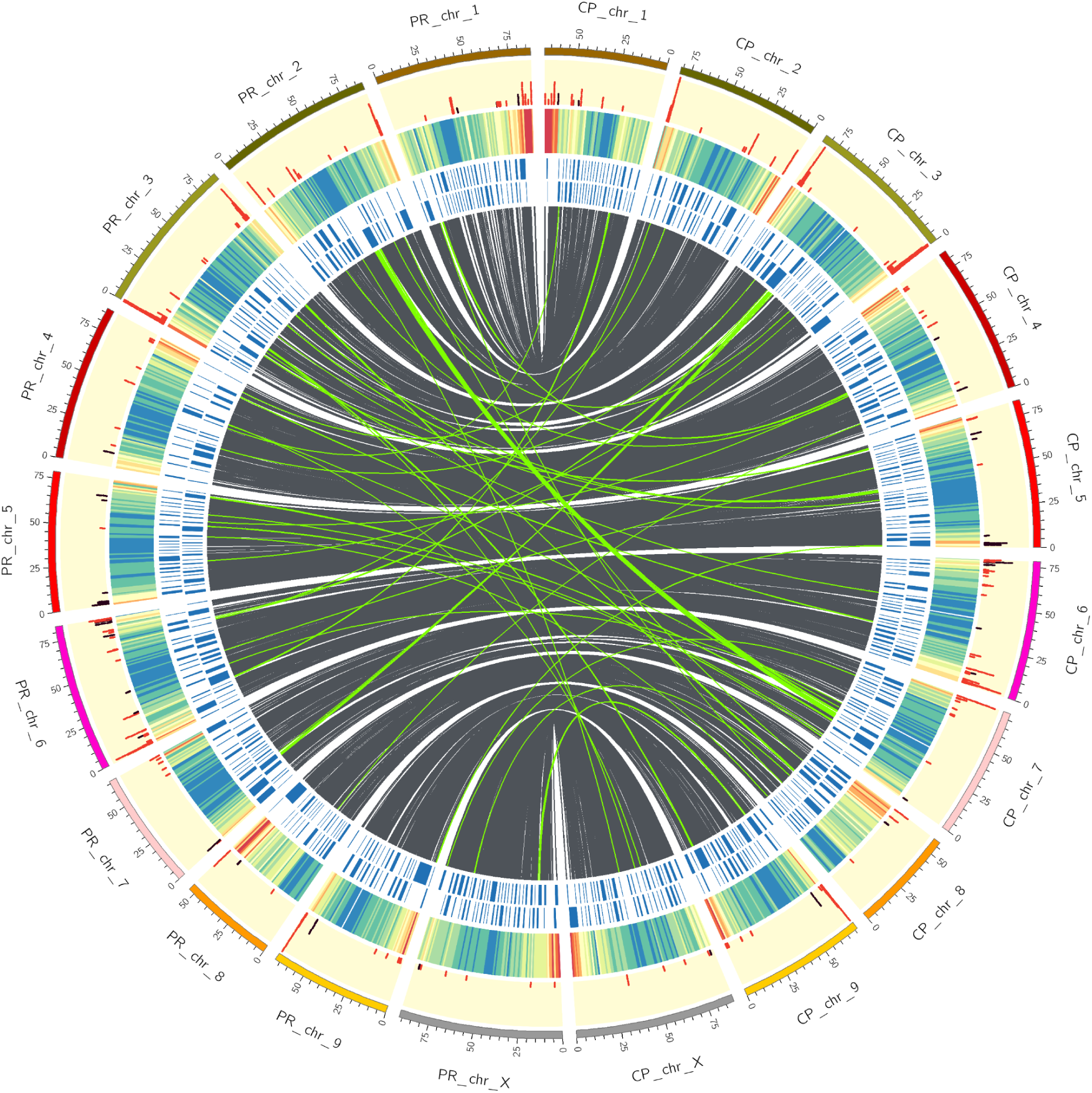
Circos plot showing, from center outwards, homologous regions (grey) and interchromosomal translocations (green), both filtered above 25kb, contig boundaries (blue), gene density (heatmap, where red is high and blue is low), and NLRs (red) and TPS (purple).

## Discussion

### HMW gDNA prep

Our method produced DNA of adequate length and sub-standard purity. Given the low yield of 34 Gb, it would be beneficial to refine the technique further, as recent reports indicate that PromethION yields of over 100Gb are now possible^98,99^. Following nuclei isolation, performing the organic extraction with phenol:chloroform^100^, in place of mere chloroform, may provide for more efficient removal of carbohydrates and proteins. As well, dark-incubation of the shoots for 3d before purification may reduce carbohydrate content^101^.

The decision to not fragment the HMW DNA surely decreased yield, due to accelerated nanopore failure when reading ultra-long fragments^102^. However, as the cannabis genome is known to be littered with repeats of 30-45 kb^9^, the 4x of ultra-long (>50kb) coverage found here is likely sufficient to resolve some of the long repeats that might falsely collapse in the absence of ultra-long coverage. Therefore, unfragmented DNA appears to be the optimal use of the ONT platform, with the caveat that sequence yield is a function of purity.

### Genome Size Estimation

Previously, flow cytometry of Cannabis nuclei has reported haploid female estimates of 818 Mb^103^ and 875 Mb^104^. Similarly, existing chromosome-scale female assemblies have total base lengths of 714^9^, 796^105^, 812^96^, and 914^106^ Mb.

From short read kmers, PR falls into this range, with estimates of 823 and 820 Mb in homozygous and heterozygous mode respectively, excluding presumed errors. CP, however, gives a size of 43 Mb (hom) and 956 Mb (het), indicating its kmer distribution is not a good fit for the model. FindGSE and other kmer-based genome size estimators suggest a minimum input of 25-30x^67,107^, with failures reported at lower coverage^108^, and so we assume that the 14.4x used here was simply inadequate.

We repeated the estimate using the binned, NECAT-corrected long reads. The homozygous estimates of 784 (PR) and 752 (CP) Mb are close to the total contig lengths of 740 and 724 Mb, indicating that this method, which has not been previously reported, appears to provide usable estimates. As these readsets should represent individual haplotypes, it is not unexpected that findGSE failed to complete in heterozygous mode.

### Assembly

#### Contiguity

Trio-binning the long reads before assembly has been shown to increase contiguity in both animals^109^ and plants^110^, but has not previously been reported for Cannabis. In this study, we follow the pattern of the original method, which includes separating reads based on parental 21-mers and discarding the unbinned. As well, we removed 21-mers with homopolymers of length 5 or greater, as these are likely to be erroneous in ONT reads^111^.

PR received about 5% more sequence than CP, and also produced a more contiguous and gene-complete draft, which suggests that the binning could have been more perfect. The switch rate of ∼3% is high compared to recent reports, based on Hi-C phasing, that return switch rates less than 1%^112–114^. These imperfections almost certainly relate to the error rate in the raw reads, which was estimated at 1.8% by comparison with short read kmers. With newer ONT reagents providing precision above 99%, the size of these error terms should naturally reduce in future.

A second experiment, in which all reads were first corrected in NECAT and then binned, produced a more equitable distribution; however, the assembly contiguity was reduced by half (data not shown). This could be the result of reads from one haplotype being corrected by the other, producing bubbles of heterozygosity which then result in excessive trimming of the corrected reads.

Additionally, given the good performance of NECAT’s error correction module, filtering reads by quality appears to not be necessary. The first phase of NECAT’s process corrects each read according to the 12 best matching reads, and then trims unusable sections^68^. This permits salvaging usable portions of long reads with low-quality regions. In the case of CP, filtering by quality at 7 prior to NECAT reduced coverage from 17x to 15x, and produced an assembly that was 1.3% shorter (740 Mb vs 749.7 Mb) and 12% less contiguous (N_50_ 1.4 vs 1.6 Mb).

After assembling the corrected reads, the raw reads are again used to form ‘bridges’ between high-quality contigs, with three reads being sufficient to establish contiguity. Because ultra-long reads are more likely to include low-quality sections^68^, not filtering them from the assembly *a priori* surely increases contiguity, at the risk of some gaps being filled with sequence of lower base-level quality.

For scaffolding to chromosome scale, ntJoin has been shown to be rapid and precise^75,115^. NECAT has been shown to have a very low rate of misassembly^116,117^, and plant genomes are known to be highly divergent^85^, so the ‘no_cut=true’ option was used in ntJoin to prevent contigs from being broken when arranged to the heterologous genotype. As well, the ‘overlap=false’ option rescues 140 BUSCO genes that were lost when ntJoin was permitted to merge contigs thought to overlap.

#### Completeness

Comparing BUSCO scores of these assemblies to previous reports highlights the value of trio binning. As shown in Table 3, both PR and CP have more single and fewer duplicate single-copy orthologs than the reference and other published assemblies, and similar numbers of fragmented and missing. By not producing alternate haplotigs, these fully-phased drafts are more frequently able to locate one SCO rather than two, which suggests that multicopy paralogs are also likely to be counted more accurately than in pseudohaploid assemblies.

#### Correctness

Yak estimates the quality of a long-read assembly by penalizing kmers found in it that are not found in a corresponding short-read dataset. Here, because the short reads are from the two parents, these figures may be somewhat inflated due to an inability to detect ONT errors that recapitulate F_0_ heterozygosity. By this metric, PR and CP have quality values (QV) of 29.2 and 28.8, in both cases corresponding to base-level accuracy of 99.9%.

While these numbers may pale in comparison to recent human^118^ and Arabidopsis^119^ assemblies, which exceed Q60 by combining ONT and PacBio HiFi datasets, they trail more typical recent plant assemblies by smaller margins. For example, a haploid banana genome assembled from 177x of ONT coverage gave a QV of 38.8^98^, and an assembly of *Ammopiptanthus nanus* from 73x of PacBio Sequel reads gave a QV of 38.95^120^. Prior to 2020, base-level accuracy of 99.9% as compared to short reads was considered top-notch^121,122^, which illustrates the rapid progress in the field.

Ultra-long assemblies frequently suffer in this metric, as they are able to assemble, perhaps somewhat erroneously, long repeats that are simply absent from other drafts. This compromise may be especially inherent in NECAT’s bridging module, which uses uncorrected long reads to bridge the highly accurate contigs assembled from error-corrected reads, and therefore will increase contiguity by introducing regions of lower quality. For both PR and CP, bridging doubled N_50_ (data not shown), and so, given NECAT’s good performance across a range of taxa^98,99,116,117^, some reduction in QV seems a tolerable trade-off.

These very good results are also attributable to rapid advances in ONT basecalling, with the raw reads in this study enjoying precision much higher than pre-2020 ONT reads and comparable to non-HiFi PacBio data. Finally, the inclusion of about 4x in raw reads longer than 50kb surely helps to span repeats that would collapse with read lengths of 15-20kb. When one compares BUSCO scores, the ability of these phased assemblies, built from 18x of ONT coverage, to outperform unphased assemblies built from ∼100x of PacBio data^96,105^ is remarkable.

#### Diploid assembly

We confirm here the results reported by Nie^81^, where trio-binning offers, by far, the best phase separation, and PECAT represents a good option for parent-naïve diploid assembly of noisy long reads. Shasta’s diploid mode, which the authors acknowledge is ‘somewhat experimental^123^,’ offers a draft with very good haplotype resolution, but its small size and low gene completeness render it sub-par for downstream analysis. We note that the readset analyzed here has N_50_ and coverage about half what is recommended. GFAse^82^, which aims to ‘unzip’ linked haplotype bubbles through the use of Hi-C contacts or, as tested here, parental kmers, does more than triple the N_50_ of the draft, but at the expense of many inter-haplotypic joins.

### Gene Predictions

#### WGS

The placement of 97.3% or 100.1% of genic reference annotations on these two drafts suggests that they are essentially gene-complete. While the cs10 reference is not competitive with new drafts produced since 2020, its annotations, which are based on several RNA-seq datasets as well as a curated set of *ab initio* predictions, provide a solid basis for annotation.

#### CN synthases

Cannabis is unique and valuable due to its synthesis of cannabinoids. The ecological function of these metabolites has not yet been proven, and here we speculate that their primary function is to influence the behaviour of mammalian seed dispersers. Other theories, such as protection from ultraviolet light^124^ or pathogens^125^, are not satisfying as they do not explain why CNs are naturally found in abundance only on female seed bracts, or why their concentration peaks after seed maturity. As well, perhaps due to their cytotoxicity^126^, CNs are synthesized almost exclusively in stalked trichomes separated from the plant body by several millimeters. Therefore, it stands to reason that their ability to protect the plant from mesophyll-feeding microbes is minimal. However, the behavioural pattern that consumption of CNs frequently induces, e.g. a period of euphoria and/or paranoia^127^, followed by hunger and/or drowsiness^128^, with a residual negative effect on short-term memory^129^, can easily be explained as a plant that would like rodents to cache more of its seeds than they might otherwise, and then forget where they put them.

The scant corroboration regarding the location of the B locus, and the number of CN synthases found in it, illustrates the difficulty of assembling this repetitive region (Table 4). Because of the importance of cannabinoids to the Cannabis space, and because of the difficulty in assembling the B locus, it offers a certain parallel with the human major histocompatibility locus, which, due to its importance to human health and ‘notoriously difficult to assemble’ structure^130^, has become a common benchmark for assembly^131^ and variant-calling^132^ tools. Here, we do not claim our results to be definitive.

All drafts place B on chr7; however, the location varies. PR and Abacus place the active synthase in the range of 50-60 Mb, in a cluster with 5 degenerate paralogs, while cs10 and Cannbio-2 place it at 30-35 Mb, in a cluster of one active and 10 inactive copies. JL recapitulates the PR/Abacus structure of one active and 5 inactive copies, but places the locus at 90 Mb. CP appears to offer a distinct arrangement, with the active synthase found in a primary cluster abutting 4 degenerate paralogs at about 62 Mb, and a secondary group of presumably inactive synthases with 88-89% identity located at 40 Mb.

Given the demonstrated difficulty of assembly, analysis of additional drafts is needed to conclusively resolve what degree of variation at B is biological and what is technical. The B_D_ and B_T_ alleles have been reported to recombine rarely, if at all^22^, despite their high homology, which may be due to one or more large SVs in and around this important locus. In other species, such as corn^133^ and sunflower^134^, massive haplotype blocks have been shown to not recombine, which prevents the separation of alleles that may provide more selective advantage as a group.

We speculate that the absence of haplotypes containing two or more putative active synthases relates to the heterozygote phenotype, which appears to occur at much higher frequency in wild cannabis than in cultivated populations. If the CBD:THC ratio of 6:5 is optimal for modulating the behaviour of seed dispersers, any imbalance in copy number might create a maladaptive divergence, and therefore be selected against. Similarly, as medical research into the therapeutic profiles of various cannabinoid ratios continues, it may be straightforward to reactivate one or more paralogs, via gene editing, to produce novel CN ratios that can be stably inherited and also customized to meet the needs of the clinic.

#### TPS

In the 1960s and 70s, Americans rapidly gained access to a wide variety of cannabis products from all over the world. Smuggling groups such as the Brotherhood of Eternal Love^135^ and the Black Tuna Gang^136^ introduced flowers and hashish from South Asia and various parts of Latin America, especially Mexico and Colombia^137^, which the growing hippie culture then propagated throughout the land.

Soon, there was a widespread perception that cannabis from different parts of the world conferred unique effects. Consumers quickly learned that *indica* types from Morocco, Afghanistan, and Pakistan offered a more relaxing, soporific, ‘body’ effect, while tropical *sativas* from Thailand and Colombia conferred a more energizing, psychedelic ‘head’ high^138^.

In the modern era, the neverending proliferation of Cannabis varieties relies, in part, upon this belief, that the consumption of different genotypes can elicit different psychoactive and somatic effects^139^. Thought to derive from synergistic interactions among cannabinoids (CNs), terpenes, and perhaps other phytoalexins, this ‘entourage effect’ has been elusive to pin down in the lab^27^.

There are at least two conceptual frameworks that may be used to dissect entourage. The more common is to view cannabinoids, such as THC or CBD, as the primary molecule, and terpenes as secondary additions that modulate the effect of the CN^140^. A second framework, proposed by David Watson of HortaPharm BV, is that the effect of THC is inherently neutral, or perhaps even ‘boring’, and that it is more useful to consider CNs as compounds that profoundly potentiate the effect of mildly psychoactive, terpene-rich essential oils.

Following many generations of wide hybridization, the landrace-based designations of *sativa* and *indica* no longer have phylogenetic value^141,142^. Yet, they are widely used in the marketplace to denote strains that have relatively more pronounced mental or physical effects^143^. Recent research has shown that these characterizations are agnostic to respect with genetic distance, but do correlate well with several terpenes: strains labelled *sativa* are more likely to be rich in bergamotene and farnesene, while *indicas* are more likely to contain myrcene, guiaol, and β- and γ-eudesmol^32^.

It may be the case that panic attacks, which are increasingly common with newly widespread use of high-THC cannabis products^144–147^, are encouraged with solvent-based processing tactics that retain most cannabinoids, but tend to lose terpenes upon solvent removal^148–150^. More speculatively, breeding strategies based on wide crosses may have created new terpene profiles not found in nature, that when ingested may contribute to a sense of unease about one’s environment.

Plants evolved terpene synthetic capacity to communicate with one another^38^, or insects^151^, or other animals^152^, to convey information about parasites^37^, food availability^153^, et cetera. Similarly to the fruit which is bitter when young, but sweet when ripe, Cannabis may speak ‘terpenese’ to advertise the presence of nutritious, fully mature seeds to its animal dispersers.

Recently, several studies^16,41,154^ have arrived at the same scheme for classifying the Cannabis population by terpene content: three groups in which the profile, or terptype^154^, is comprised primarily of myrcene (MYR), terpinolene (TER), or limonene with caryophyllene (LIM).

Because these and other terpenes frequently show anxiolytic^155–157^ and antidepressant^158^ effects in animal^159,160^ and human^161,162^ models, understanding the genetic basis for their accumulation in Cannabis is valuable, for both the clinic and the dispensary.

Previously, the products of 33 Cannabis terpene synthases had been quantified via heterologous expression in *E.coli*^88^. Via BLASTx, we were able to verify many of our predictions, clarify others, and also identify possible lacunae to be resolved in future investigations.

Based on field observations, which corroborate the grey literature, we postulate that PR has a LIM terptype^163^, and CP a MYR terptype^164^. Therefore, it is notable that XP_030500628.1, a gene predicted to encode ‘(-)-limonene synthase, chloroplastic like’, has its best (99.8%) hit in cs10 and CP to CsTPS14: Canna Tsu (-)-Limonene, while in PR it is (99.2%) to CsTPS1: Skunk (-)-limonene. The difference is small, yet cursory evaluation of an alignment reveals, among other polymorphisms, a proline-serine transversion between the two groups, which indicates that the alleles are in fact distinct (Fig. 7). We also note that XP_030501051.1, a myrcene synthase, is confirmed with best hits to CsTPS15: Canna Tsu Myrcene in PR, CP, and cs10. However, the Grade in cs10 and CP is quite good (96.6% & 96.7%), while in PR it is much lower (75.7%), with a CDS that includes several premature stop codons (Fig. 8).

While admittedly scant, these data suggest that the difference between the MYR and LIM terptypes might derive from different numbers of functional myrcene synthases. Because these myrcene and limonene alleles lie within the same MTC on chromosome 5, and gene clusters such as these are frequently co-regulated^165^, analisis of the sequence variation that lies within them is worthy of further inquiry.

On chr1, we found that XP_030491253.1, a predicted (3S,6E)-nerolidol synthase 1, returned a very good (99.5%) hit to CsTPS35: Lemon Skunk Linalool/Nerolidol, a TPS that is among the most highly expressed in several cultivars, and which *in vitro* produces 95% nerolidol from farnesyl pyrophosphate (FPP), but 93% linalool when fed geranyl pyrophosphate (GPP)^88^. In Cannabis trichomes, GPP is more abundant than FPP^13^, and linalool is about as common as nerolidol^154^. Both have been shown to exert anxiolytic effects in animal models^166,167^. Therefore, we imagine that these nerolidol synthase annotations may be meaningful when seeking to evaluate genetic variance in linalool content.

We were also able to clarify the role of XP_030484762.1, ‘probable terpene synthase 9’, which produced a perfect hit to CsTPS29: Blue Cheese Linalool in cs10, and 99.2% hits in PR and CP. Similarly, the 4 TPS on chr9, all predicted as ‘probable monoterpene synthase MTS1, chloroplastic’, gave near perfect hits to proteins demonstrated to produce primarily myrcene, terpinolene, or a mix of geraniol and himachalene (Table 8).

Several loci did not find good matches among the characterized enzymes. In particular, the (E,E)-geranyllinalool synthases on chr7 and many of the diterpene synthases on chr1 and chr6 had best hits with Grade <80%. Expression *in vitro* of these types could provide a fuller picture of the terpenes that contribute to entourage. For the future, we hope to characterize the three MTC as polygenic Mendelian units, which are likely to be the major contributors to genetic variance in terptype. Because of their position in subtelomeric regions (Fig. 11), they are much more likely to be affected by recombination^9^, and so may represent some of the fastest evolving regions of the Cannabis genome.

#### NLRs

In rosids, the number of NLRs ranges from 58 to 930^168^, and so the counts reported here (227 and 240) are not atypical.

Clustering of NLRs is consistent with the theory that their diversification results from duplication via unequal crossing over, followed by neofunctionalization^169^. And, the placement of several clusters at the very ends of chromosomes is consistent with their rapid evolution^170^, especially given that in Cannabis, large central portions of chromosome bodies appear to be insulated against recombination, with most events restricted to their distal ends^22^.

NLRs are notoriously difficult to assemble from short reads^171^, in part due to strong conservation in the NBS domain, with collapsed paralogs and technical chimeras being common artifacts. Sequencing reads of sufficient length to span one or more full-length genes offer more clarity as to cluster structure^56^, with phasing of haplotypes offering further improvements^172^.

This increased resolution becomes meaningful as trait mapping commences for Cannabis. At present, there is only one R-gene reported, PM1, which confers qualitative dominant resistance to the powdery mildew pathogen *Golovinomyces ambrosiae*^60^. While the PR x CP F_1_ has been observed to be susceptible to powdery mildew, accurately assembling this cluster in resistant genotypes will likely be the most efficient path towards elucidating the biochemistry of perception.

Linkage mapping places PM1 in MRC2b, which contains 16 NLRs in PR and 18 in CP (Table 7). PR assembles the cluster in one contig while CP divides it among three. Visualizing the lifted-over reference annotations, the Cannabis-specific NBS HMM hits, and the NLR-Annotator predictions illustrates the convergence and divergence among callsets (Fig. 12).

**Figure 12.**
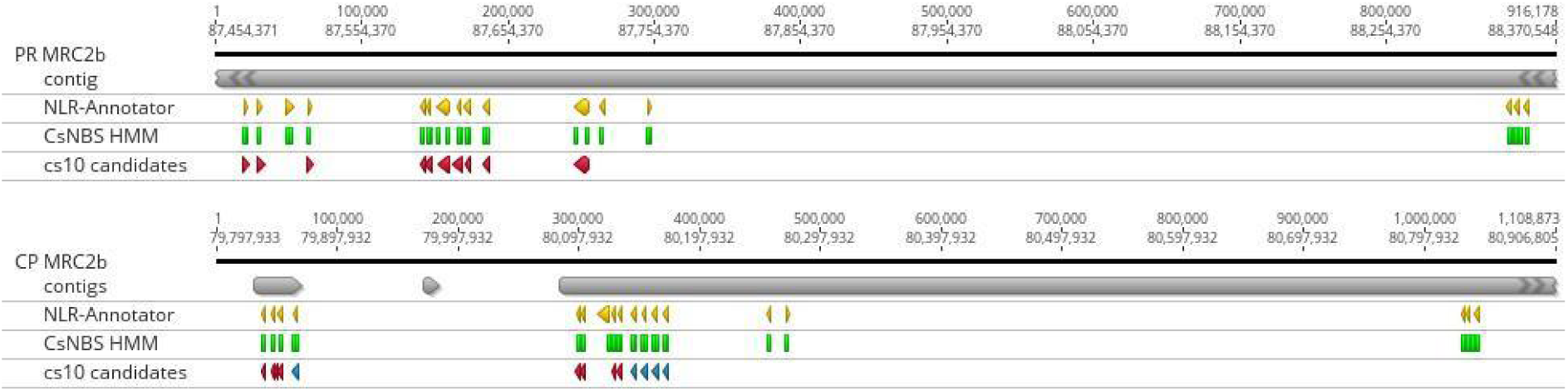
NLR predictions from NLR-Annotator (yellow), the CsNBS HMM (green), and the cs10 reference (red) found in the contigs (grey) that constitute MRC2b in Punto Rojo (top) and Cherry Pie (bottom). In the cs10 track for CP, the 5 homologs of XM030648577.1 are marked in blue, and the best match is at left.

The reference predicts 10 NLRs in this region of about 1.5 Mb. In PR, all were present in single copy. In CP, 8 are present as a single copy; one, XM030647777.1, is absent, while another, XM030648577.1, has four additional copies with exonic identity over 99%. The ordering of these genes varies among PR and CP and cs10. In both PR and CP, the HMM and NLR-Annotator both predict one additional NLR within the canonical cluster, and 5 additional candidates in the ∼650 kb downstream.

Throughout the genome, we observed that NLR-Annotator made about 40 predictions, mostly under 1kb, that were not corroborated by the HMM or the cs10 annotations, which we presume to be false positives. The HMM had only a few hits that were not corroborated, but sometimes finds two hits in one gene, particularly on chr2. Therefore, the intersection of the two methods was taken as a parsimonious set.

#### Comparative Genomics

At the level of contigs, these results compare favorably to recent assemblies with much higher coverage.

CP and cs10, the NCBI reference, are thought to be related as both are CBD clones in the ‘Cherry’ family which arose in Colorado following legalization in 2012. cs10 gDNA was fragmented to 15kb, sequenced on the ONT platform to a depth of 100x, basecalled with Guppy 3, assembled in miniasm, polished with Racon-Medaka and Pilon, scaffolded with a Hi-C library, and super-scaffolded at chromosome scale using a linkage map derived from an unrelated (Skunk x Carmen) F2 population^9^.

The contig number, N_50_, and total length of CP are rather similar to cs10, which suggests that longer length and higher accuracy, plus trio-binning, can effectively compensate for lower coverage. In particular, the higher accuracy of Guppy 5 and the good performance of the NECAT assembler, perhaps especially in the error correction phase, appear to allow confident assembly through many repetitive regions with as little as 5x of coverage. The cannabis genome is known to be littered with repeats of 30-45kb^9^, and so the similar N_50_ and contig number may indicate common zones of difficulty that may require additional effort to resolve.

To scaffold the contigs to chromosome scale, a collection of superscaffolds from the recent Salk Institute data release was chosen (Table 1). These assemblies are assembled from PacBio HiFi reads and scaffolded and phased with Hi-C libraries^76^. We observed many fewer small inversions than when scaffolding to other recent long-read assemblies (data not shown), which likely reflects an enhanced ability of Hi-C to properly orient contigs when leveraged against the greater accuracy of HiFi reads. Scaffolding to any one haplotype invariably produced several troubling large-scale rearrangements, and so a collection of chromosomes was chosen that produced a visually acceptable dotplot.

When PR and CP are aligned to one another (Fig. 10), large inversions can be seen on chromosomes 3, 5, 7, and 9, which are absent when each is aligned to its substrate (Fig. 9). These large inversions may inhibit recombination, as has recently been shown for tomato^99^.

We note that, at the level of SVs, PR appears to be more diverged from its substrate than CP: it shows less synteny, (538 vs 679 Mb) and more inversions (8.23 vs 2.05 Mb), translocations (35.9 vs 19.8 Mb), and unaligned regions (206 vs 68 Mb). Meanwhile, the number of SNPs varies much less (2.37M vs 2.27M), highlighting the value of counting larger variants (Table 10). While scant, these observations suggest that Punto Rojo, a long-flowering landrace of only moderate cannabinoid content, may represent an unusual lineage that remains undersampled among the current crop of Cannabis genomic resources. A large number of anecdotal reports suggest that Colombian landraces were a common founder of modern drug types^173^, and so future work should include the sequencing of more Colombian heirlooms, in order to identify characteristic genes or haploblocks that may have persisted in the modern market.

It has been shown that SVs are called more accurately from *de novo* assemblies than from mapping long reads to a reference, especially for variants over 100kb^174^. SyRI, one of the few tools capable of such an analysis, here shows that PR and CP share 505 Mb (66.5%) of synteny, and have 96 Mb of detectable inversions, duplications, and translocations, leaving 206 Mb (27.1%) unalignable. This may seem imprecise when compared to opisthokont genomes that routinely show synteny above 90%^85^, but more likely reflects the greater intraspecific architectural diversity found in plant genomes, which has only recently become quantifiable, with benefit of third generation sequencing. For comparison, when aligning two gold-standard maize genomes (PH207 and B73), SyRI found 62.2% synteny and 32.5% unalignable^85^. It may be that anemophilous outcrossers are particularly unlikely to purge rare variants, and so in future we hope that the creation of a Cannabis pan-genome can further characterize the structural variation that exists across its range.

## Conclusions

Here, we show that trio-binning can separate noisy ONT R9 reads, and produces very good fully-phased assemblies. By avoiding haplotype collapse, we are better able to characterize the content of two important gene classes, which occur in clusters of paralogs, and represent the fastest-evolving regions of the Cannabis genome. We are also able to ascertain the presence of many large structural variants, which are frequently invisible when mapping to a reference.

The natural diversity of Cannabis is remarkable; few species can be found from zero to sixty degrees of latitude and at altitudes from 0-3000m. Further characterization, including additional genome assemblies and especially multiple genotype, multiple environment field trials, should enlighten as to the variants that facilitate adaptacion to such a wide range of habitat.

The PR and CP parents have both been used to create a wide variety of testcrosses, and we hope that these new assemblies will enable more precise trait mapping than would be possible with an exogenous reference.

## Declarations

## Acknowledgements

We thank the farm team of Medcann Colombia for plant care and for producing the initial cross of PR x CP.

We thank Ruta Sahasrudbe and Lutz Froenicke for preparing and sequencing the genomic libraries. The sequencing was carried out at the DNA Technologies and Expression Analysis Cores at the UC Davis Genome Center, supported by NIH Shared Instrumentation Grant 1S10OD010786-01.

We thank Juan Guillermo Torres Hurtado for facilitating the use of the Pontificia Universidad Javeriana’s ZINE high-performance computing cluster.

## Availability of data and materials

The genomes are available from NCBI under accession codes JBDLLE000000000 (Punto Rojo) and JBDLLD000000000 (Cherry Pie). Additionally, copies named according to PanSN-spec^175^, with annotation GFFs, are available from Zenodo (DOI: 10.5281/zenodo.15284085).

## Authors’ contributions

BP conceived and designed the project, performed the bench work, assembled and annotated and compared the genomes, and drafted the manuscript. AK advised on various aspects of genome assembly and annotation, and developed the Cannabis-specific HMM and the core files for the Circos plot. WT supervised the project and edited the manuscript.

## Ethics approval and consent to participate

Not applicable.

## Consent for publication

Not applicable.

